# A Superior “Thumb” Drive: Optimizing DNA Stability for DNA Data Storage

**DOI:** 10.1101/2023.05.11.540302

**Authors:** Jiun-You Liu

## Abstract

DNA digital data storage is an emerging technology with the potential to keep up with increasing demands in space, energy, and materials used for information storage. To make DNA storage systems more practical and cost-effective, it is crucial to determine how the physical and chemical properties of DNA affect its performance. Thus far, scientists have experimented with various DNA storage mediums and encoding techniques to maintain DNA stability. In this study, we first review existing strategies for maintaining DNA stability under various environmental constraints, organized according to the application of the storage system (Archival Storage, Working Storage, and Dynamic Storage). Then, we report on our own quantitative evaluation of the tradeoffs between storage density and DNA stability to identify the optimal system configurations for distinct storage scenarios.

## Introduction

The digital revolution has heralded an era of unparalleled connectivity and information exchange, transforming the way we live, work and interact with one another. Data is at the heart of this transformation, driving numerous aspects of our lives, from the proliferation of social media and e-commerce to the rise of automation and artificial intelligence. These data-driven technologies have led to an explosion of generated data. In fact, the total amount of data created and stored worldwide in 2018 was a staggering 33 zettabytes, the equivalent of 66 trillion hours of music or 1.2 million years of high-definition video.^1^ By 2020, this number had skyrocketed to 59 zettabytes, with projections forecasting a mind-boggling 175 zettabytes by 2025.^2^

With information being produced and consumed at an unprecedented rate, traditional storage systems are struggling to keep up with the demand. Hard drives and flash drives have limited storage capacity and become corrupt over time. Meanwhile, cloud-based storage solutions that house internet data in remote data centers have become increasingly energy-intensive and expensive to maintain.^3^ If we continue to consume storage at this rate, we will face two significant problems by 2025. Firstly, the amount of readily available high-grade silicon, a natural resource used to manufacture memory chips, will not be enough to keep up with the amount of data being generated. Secondly, data centers are projected to consume around 20 percent of global electricity production, leading to a massive carbon footprint predicted to match that of the transportation industry.^4^

These challenges highlight the urgent need for innovative and sustainable storage solutions. One such emerging technology that has seen numerous advancements in recent years is DNA data storage. DNA, the genetic material found in all living organisms, is compact and durable, making it an ideal candidate for efficient and reliable data storage. DNA molecules can store information using the four nucleotide bases: Adenine (A), Thymine (T), Cytosine (C), and Guanine (G). These four bases serve as the building blocks of our genetic information and can be encoded to represent digital information. This encoding process involves converting digital data into binary code (zeroes and ones), then mapping them onto the four nucleotides (A, C, T, and G). The resulting DNA sequence can then be synthesized in the lab and stored for extended periods of time.

The most significant advantage of using DNA storage is its exceptional storage density. Due to the minuscule size of DNA molecules, they can store an enormous amount of data in a very compact space. One solitary gram of DNA has the potential to store up to 215 million gigabytes, or 2.15*E*^−4^ zettabytes of data, making it the most space-efficient storage solution to date.^5^ To put this into perspective, using DNA storage technology, all the data ever recorded in a container as small as a shoebox.^6^ This is in stark contrast to the world’s most advanced USB thumb drive, which can only store up to one terabyte, or 1. 0*E*^−9^ zettabytes of data^7^. Now, if we calculate the amount of DNA present in our biological thumb, there is around 100 million times that storage, which makes DNA a far superior “thumb” drive.

Another substantial advantage of DNA storage is its energy efficiency. Once the DNA has been synthesized and the data written onto it, the information can remain stable with minimal energy input. In contrast, traditional storage solutions require a constant energy supply to maintain data integrity. For example, a hard disk needs a steady supply of electricity to power the spinning disk and the read/write head that accesses the data. Traditional storage solutions also require energy to maintain a cool environment to prevent overheating.^8^ DNA storage does not require any cooling, as DNA molecules can be stored at room temperature with a marginal impact on their stability. Thus, DNA storage can potentially reduce energy consumption significantly and help organizations meet their sustainability goals.

Despite offering several advantages over traditional storage solutions, DNA storage has numerous limitations, one of the most significant being DNA stability. DNA is susceptible to damage from environmental factors such as UV radiation, high temperatures, humidity, oxidative stress, etc., all of which cause strand breaks or mutations in the DNA molecules, leading to errors in the stored data or even complete loss of the information.^9^ To resolve these challenges, researchers have developed strategies such as redundant synthesis and error correction codes (ECCs) to improve the stability and reliability of DNA storage systems.^10^ Redundant synthesis is a strategy for improving the reliability of DNA data storage by synthesizing multiple copies of the same data in different DNA molecules, which are then stored in parallel. This approach reduces the risk of errors or damage to data integrity, as even when a single copy of the data is damaged or contains errors, the redundant copies can be used to reconstruct the original data with high degrees of accuracy. Like electronic storage systems, error correction coles are another strategy used in DNA data storage to improve reliability. ECCs add additional information to the DNA sequence in the form of redundant bits, or overheads, to detect and correct errors in data storage. For instance, a simple ECC might involve adding a parity bit to each nucleotide, which can be used to correct single-bit errors in the nucleotide sequence. These codes introduce sufficient redundancy to mathematically recalculate and recover the original data, even when errors or lost strands exist.

Though redundant synthesis and ECCs are effective strategies for improving storage system reliability, they both involve synthesizing additional redundant DNA sequences in advance to account for potential errors. This leads to a tradeoff between storage density and DNA stability: the more redundancy and error correction overheads are used, the less digital information can be stored within each DNA strand. This tradeoff between error correction and information density must be optimally balanced to achieve the most effective storage system. The balance must be carefully considered based on the specific use case of the storage system since DNA stability can fluctuate depending on the storage environment and how it’s operated. For example, a system that is expected to be exposed to high radiation levels or other forms of damage may require more redundant sequences and error correction overheads to ensure the data’s integrity. In contrast, a controlled environment with minimal potential damage factors would allow the system to prioritize information density over error correction.

In recent years, scientists have developed various DNA storage types to address this tradeoff. These systems are tailored to specific storage applications, such as long-term Archival Storage and frequently-accessed Working Storage, with their configurations optimized to meet the unique requirements of each application.^11^ This classification enables researchers to manage the tradeoff better because the more frequently stored DNA is accessed, the higher the probability of physical damage to the DNA strands, such as strand breaks, chemical modifications, and oxidation, which all result in DNA degradation and loss of data. For instance, Archival Storage is designed for long-term data preservation and does not require frequent access to the stored data. As a result, it can prioritize high storage density and long-term stability over its reading and writing speeds, necessitating fewer redundant overheads. On the other hand, Working Storage is designed for applications that require more frequent access to data, such as cloud storage and data centers. As a result, it is optimized for high read and write speeds and can handle numerous read and write cycles; however, it also has a much lower storage density than archival storage due to its high redundancy requirement.

To explore the possibility of making these DNA storage systems more practical and cost-effective, it is crucial to determine how the physical and chemical properties of DNA affect its performance as a storage medium. Since DNA is the building block of these storage systems, helping DNA maintain its stability while it undergoes processing strains and environmental conditions will be a major design challenge. Thus far, the majority of researchers have attempted to improve DNA storage by creating novel DNA storage mediums or encoding techniques; however, there is a research gap in examining how efforts to maintain DNA stability can impact information density. Specifically, there is limited research pertaining to how certain system configurations affect the stability-density tradeoff.^12^ In this paper, we first review existing strategies for maintaining DNA stability under various environmental conditions, organized according to the application of the storage system (Archival Storage, Working Storage, and Short-Term Storage). Then, we report on our own quantitative simulation and evaluation of the tradeoffs between storage density and DNA stability to identify the specific system configurations that impact this tradeoff.

## Literature Review

A generic DNA storage schema with three different configurations that each address a type of storage application is illustrated in Figure 1. Depending on the storage application ranging from Archival Storage to Short-Term Storage, the DNA in the system can undergo various forms of manipulation, all of which can severely impair DNA stability and its ability to perform as an effective storage system. To better understand how these molecular mechanisms can impact a DNA storage system, it’s important to examine its molecular architecture first. A DNA storage system consists of multiple files, each containing numerous DNA strands that are typically ∼150-200 nucleotides long.^13^ All DNA strands within a file have a shared address sequence positioned on either end of the strand, which serves as markers for the specific file. This marker can be accessed and recovered using PCR, a laboratory technique to amplify a DNA strand to preserve or manipulate genetic information. Mutations or losing a DNA strand within a file can cause loss of information or potential decoding errors. Thus, it is imperative to understand the effects of these molecular mechanisms of damage as they present opportunities for degradation that may influence the overall storage density and access frequency of the system.

**Figure 1.**
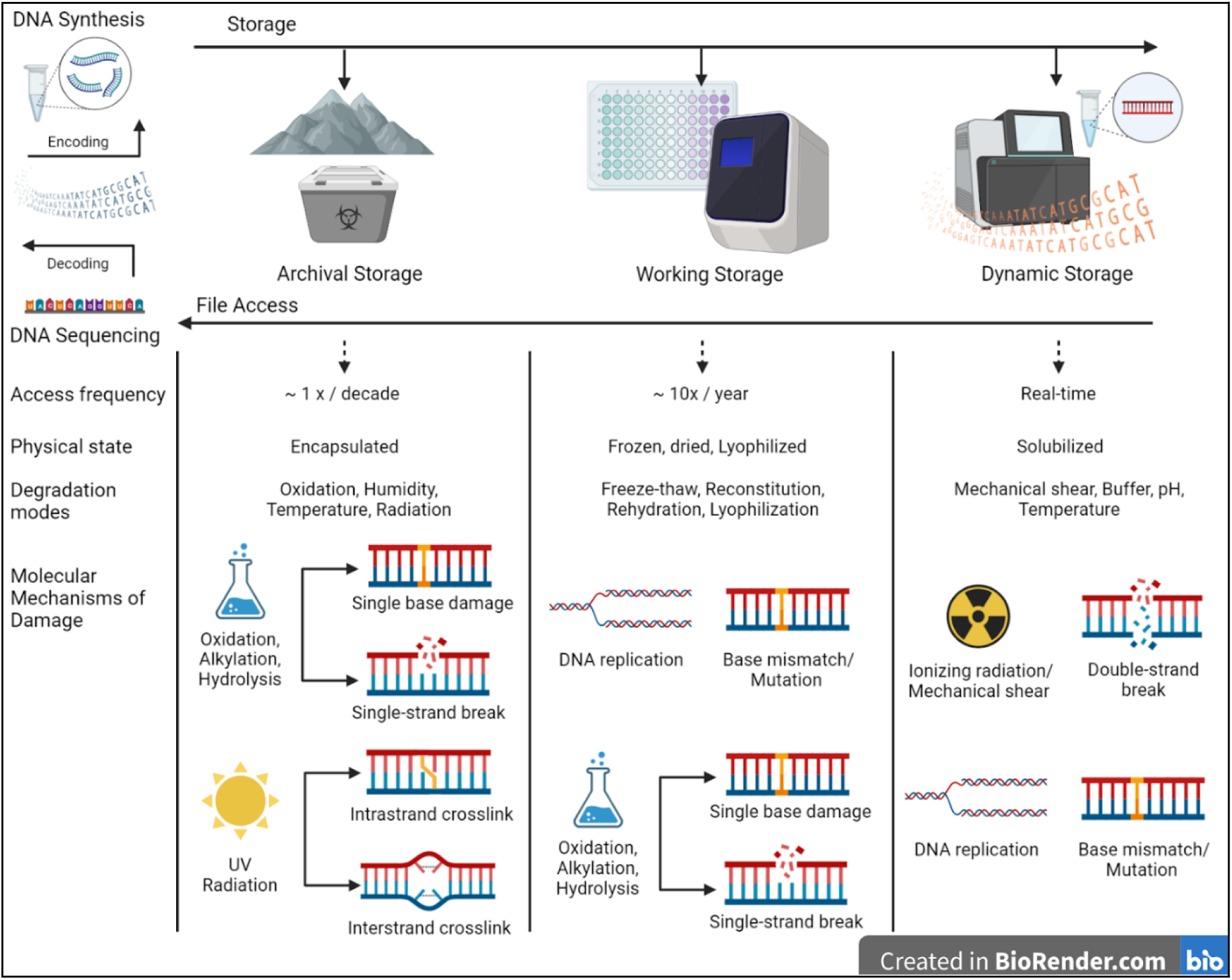
A Generic DNA storage schema showing various configurations that address diverse storage applications; The features of each storage configuration are described, along with the molecular mechanisms of damage that pertain to each configuration. Figure created with BioRender.com.^14^ ^15^ ^16^

## Archival Storage Stability Review

Long-term Archival Storage is expected to be one of the initial applications of DNA-based information storage, specifically for preserving historical records over extended periods ranging from decades to centuries. The high cost associated with synthesizing DNA is more easily justified over extended periods; therefore, this technology will likely be applied to address those storage needs.^17^ To ensure the effectiveness of DNA-based storage for these applications, it is crucial to comprehend DNA’s long-term stability.

Many researchers have pointed to the successful recovery of DNA from fossils dating back millions of years and microbes trapped in permafrost as proof of DNA’s long-term stability.^18^ However, there are several limitations to consider. One major concern is the possibility of contamination by modern bacteria or human DNA, which could lead to misinterpretation of the findings. As a result, according to theoretical calculations and physical measurements, the maximum age for DNA recovery from natural samples is estimated to be approximately 400,000 years.^19^ Next, relying on DNA stability estimations based on natural samples might lead us to assume the capabilities of DNA storage overconfidently. DNA extracted from fossils rely on ribosomal and mitochondrial DNA, of which hundreds of copies exist per cell. While multi-copy DNA strands are easier to retrieve than genomic DNA sequences, they are less informative for genetic analysis. Even when DNA is extracted from fossilized or permafrost samples, recovery is still limited to the most prevalent DNA strands because of significant degradation. This idea is reinforced by the estimation that fossilized DNA has an anticipated half-life of fewer than 500 years, causing a breakdown rate of 5.50*E*^−6^ *nucleotide*^−6^ *year* ^−1^.^20^ This fragmentation rate is significantly less than the 400,000-year expected period for DNA recovery. Therefore, the data from fossilized DNA suggests that DNA storage applications may only remain stable for a few hundred years or less.

Fortunately, engineered DNA storage systems can offer increased stability compared to naturally occurring samples by utilizing precisely regulated materials and surroundings. Various methods involving DNA in dehydrated forms have been designed to reduce the probability of breaking down its phosphate backbone.^21^ For instance, DNA can be stored as a lyophilized powder, attached to Flinders Technology Associate (FTA) paper through adsorption, embedded into silica beads, or kept in storage mediums made of biopolymers, like the commercial item DNA Stable.^22^ In some cases, the DNA is stabilized by binding onto a matrix, a combination of various biomolecules that provides a three-dimensional structure for the DNA to attach to and form stable complexes. Silk, for instance, has been found to increase DNA stability, with 80% of embedded DNA recoverable after 40 days at 25, 37, and 45 degrees Celsius, compared to only 20% without protection.^23^ Salts have also demonstrated the ability to stabilize dried DNA, especially in the presence of high ambient humidity, and to sustain high DNA loading (>20 wt%, the amount of DNA that can be incorporated into a carrier material) while still maintaining a high level of DNA accessibility.^24^

Numerous approaches have been tested to increase the stability of DNA storage systems, but the process of enclosing DNA within a non-organic substance composed of iron oxide or silica was identified as the state-of-the-art method that reliably leads to the highest DNA stability. Encapsulation has been demonstrated by numerous research groups to significantly improve DNA stability.^25 26 27^ An example is the study by Puddu et al., in which the DNA stability under unprotected conditions was compared to encapsulated DNA at a temperature of 100 degrees Celsius for half an hour. Subsequently, only 0.05% of the dehydrated, unprotected DNA remained, while 90% of the encapsulated DNA was retrieved.^28^ According to Grass et al., silica particle encapsulation could preserve DNA for 20-90 years at ambient temperatures, 2000 years at 9.4 degrees Celsius, and more than 1.5 million years at temperatures less than –20 degrees Celsius.^29^ The figures were derived using accelerated aging models that assessed DNA records exposed to elevated temperatures ranging from 65-75 degrees Celsius. Following accelerated aging, the DNA strands in the experiment were decoded back into digital information, providing evidence that entire files could be recovered.^30^ In several instances, data could be retrieved repeatedly, and only minimal degradation was observed.

Prior research strongly supports DNA’s long-term stability, especially when protected by silica. However, there are some limitations to this research. For one, many studies rely on accelerated aging models that can predict stability for millions of years using extrapolations, which as with all simulations, means that the data is at risk of deviating from legitimate stabilities as more time passes. This is especially relevant if an undiscovered degradation process exists that does not follow the typical temperature-dependent decay curve. To improve confidence in stability predictions, additional research should examine the effects of various factors that can cause DNA degradation over time, including exposure to pH, radiation, humidity, and mechanical stresses such as freeze-thaw cycles and liquid handling. To develop a better understanding of DNA stability, it is necessary to evaluate a wider range of storage conditions, including models that simulate accelerated aging under different environmental conditions. This is important for developing optimal DNA storage systems that can preserve data accurately over the long term.^31^

Accelerated aging research poses another challenge, as the techniques employed to evaluate DNA degradation are insufficiently accurate and rely on amplification steps like PCR, which may distort or influence the results. New techniques of sequencing such as deep sequencing could be employed to detect rare degradation incidents.^32^ Another potential solution is to conduct long-term non-accelerated investigations into DNA stability, lasting at least one human generation. We can then compare these real-time results to those obtained from accelerated degradation modeling, revising and enhancing the predictive models established previously.

## Working Storage Stability Review

While methods to achieve DNA stability for hundreds or even millennia are possible, they call for completely enclosing DNA inside a silica matrix. This is because a matrix shields the DNA from contact with moisture, radiation, temperature changes, and other possible reactants. It may also be the most reliable and cost-effective way of storing, requiring the least amount of specialized apparatus like humidity and refrigeration that must be closely regulated. However, this type of storage is most suitable for scenarios in which access to the DNA is infrequent or unnecessary due to the substantial DNA loss that occurs during recovery from the encapsulated medium and the more time-consuming encapsulating and extracting procedures. A Working Storage system can employ encapsulated DNA as the primary copy of data in a DNA storage system, while using less dependable but more accessible techniques to store “working” versions of the data.

Three potential storage methods for Working Storage are: DNA immobilized in an aqueous solution at temperatures of 20 or 80 degrees Celsius, refrigerated in an aqueous solution, or in a dehydrated solid state. Although all three methods have demonstrated near-100% recovery rates after two years of storage, studies involving accelerated aging at higher temperatures have suggested that these storage methods may not provide sufficient stability for several decades.^33 34 35^ For instance, one study found that when exposed to 70 degrees Celsius, dried, adsorbed, and aqueous DNA experienced notable degradation, with only 15 to 70% integrity remaining after 120 hours. In contrast, encapsulated DNA exhibited no noticeable degradation after a week.^36^

Another method that can be used to store DNA for intermediate timescales is dehydration. Research has shown that vacuum-dried DNA can remain stable for up to 22 months at a temperature of 5 degrees Celsius.^37^ Although there have been some studies on the effectiveness of recovering dried DNA, more comprehensive studies on repeated rehydration are needed. One study dehydrated a collection obecausecontaining approximately 20 kilobytes of data using a microfluidic device called “PurpleDrop.” The PurpleDrop is a small device that uses manipulable droplets of liquid to study a variety of liquid-based phenomena such as chemical reactions, biological processes, and fluid dynamics.^38^ In this case, PurpleDrop rehydration was carried out using water for varying lengths of time, ranging from 1 to 2 minutes. The amount of recovered DNA generally increased with each period, with about 77% of the DNA by mass being recovered in just 1 second. The complete file was recovered and decoded in this study because of adequate physical redundancy, a process in which multiple copies of the same data are stored on different DNA molecules, which are then stored in separate locations. This redundancy can help mitigate the effects of errors or failures that may occur during the storage process. However, it is important to note that in the researcher’s study, the retrieval process’s consistency showed significant variability, indicating that further research is needed to understand the mechanisms behind the dehydration process.

Additional forms of DNA instability include depurination and oxidation. Depurination involves the loss of Guanine or Adenine nucleotides from the DNA backbone. The speed at which depurination and oxidation take place is much higher than the strand breakage rate, with approximately 5 percent of 200-nucleotide strands depurinating each year at room temperature.^39^ The degradation of DNA can significantly impact the efficiency of storage systems. Oxidation, for instance, can create bonds between DNA strands, thus preventing the rehydration of dried DNA and the amplification of DNA. Depurination on the other hand can cause sequencing errors or missing nucleotides. Depending on the storage system’s physical redundancy and encoding methods, the expected lifespan of storage techniques, such as solution, frozen, or dried storage, may be further reduced.

## Short-term Storage Stability Review

Over a decade before the creation of the first DNA storage devices, DNA-based processing was being developed.^40^ By combining the two fields, researchers can conduct in-storage computation, which offers the potential for direct searching and editing of DNA databases.^41 42^ The ultimate goal for DNA storage systems is to operate with significantly lower latencies than the current near multi-day process of receiving and storing data, but both use cases will require actual manipulation of DNA. The intricacy of the storage technique is a key consideration for Short-term Storage. DNA for these uses will likely need to be kept in an aqueous solution that is both soluble and contains buffers suitable for molecular processes such as polymerization, enzyme-driven reactions, and transcription. Improving DNA stability in this particular setting will impact not only the operational lifespan of these systems but also the kinds of encodings that will be required.

Manipulating DNA often involves liquid handling using microfluidic channels, tubing or pipette tips. These procedures can subject DNA to shear forces, which causes strand breaks. Numerous investigations have evaluated how vortexing or other shearing tools can affect DNA integrity and have found significant deterioration, such as nearly 70% fragmentation after 3 minutes on a typical tabletop vortexer operating at maximal speed.^43^ While it is unlikely that DNA storage systems will utilize shearing devices, their use in regulated environments can help establish correlations between shear forces and fragmentation rates. Perhaps what’s more important is evaluating DNA fragmentation due to physical pipetting. A study using a 1 mL pipetting tip found that a single quick pipetting motion resulted in 70% of the lengthy lambda phage DNA being broken. Even with slower and gentler pipetting, over 50% of the lambda DNA was still broken.^44^ Given that liquid handling frequently employs even tinier 200, 20, and 2 microliter tips, which could produce greater shear pressures, this is somewhat worrisome. The pipetting actions could be considerably slowed down with automation, but this could come at a significant cost in terms of lengthening the time needed to carry out each operation or manipulation.^45^ No matter which liquid handling technique is used, it is essential to conduct a comprehensive DNA stability evaluation. Furthermore, it is crucial to explore innovative liquid handling methods while quantitatively assessing their impact on DNA stability. For example, managing liquid droplets ranging from nano to microliter sizes may lead to different effects on DNA stability due to additional pressures, such as exposure to surface tensions that arise from a high surface-to-volume ratio.^46^

## Methodology, Results, and Analysis

Up until now, this study has examined existing strategies for maintaining DNA stability under various environmental conditions, illuminating a research gap in examining the tradeoff between DNA stability and information density. To fill this research gap, we now report on our own quantitative simulation of the tradeoffs between storage density and DNA stability to identify the specific system configurations that impact this tradeoff. Simulation research provides a means to study complex phenomena or systems that may be challenging or impractical to investigate through physical experimentation alone. Creating a computational model that mimics the behavior or characteristics of the real-world DNA storage system under investigation allows researchers to study and analyze the system in a controlled and reproducible environment. To that end, we generated a simulated environment modeled after Grass et al.’s accelerated aging model to evaluate the DNA system configurations that affect the stability-information density tradeoff. We assessed this tradeoff through a series of analyses illustrated in Figures 2 to 4 below. Specifically, we demonstrate how the connection among strand loss, information density, copy number, and strand length can be mathematically represented and leveraged to design better systems.

**Figure 2.**
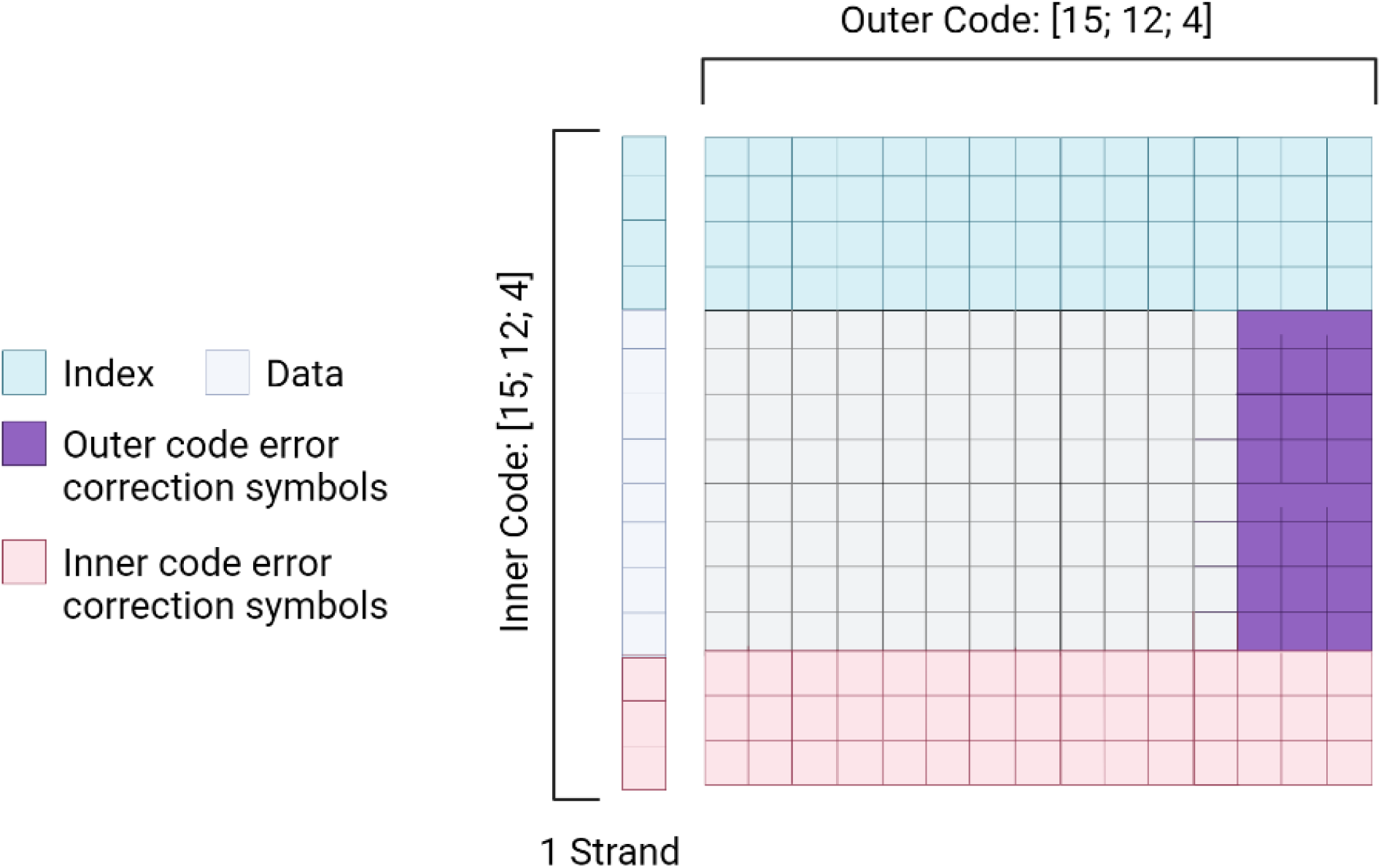
Reed-Solomon encoding scheme. Figure was created with BioRender.com.^50^

**Figure 3.**
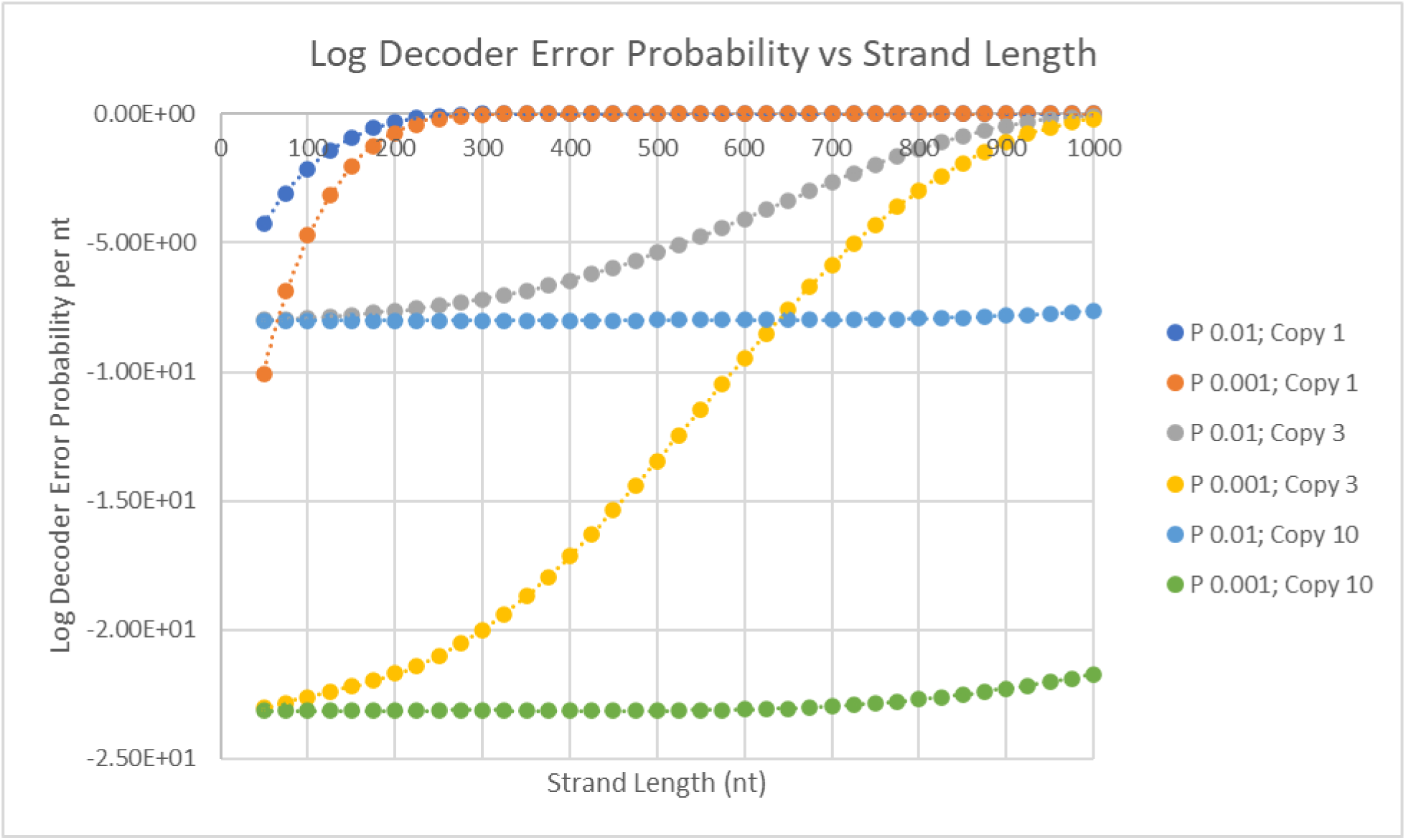
Effect of DNA strand length, strand error rates, and copy number on logarithmic decoding error probability; strand error rate is defined as insertions, mutations, and deletions.

**Figure 4.**
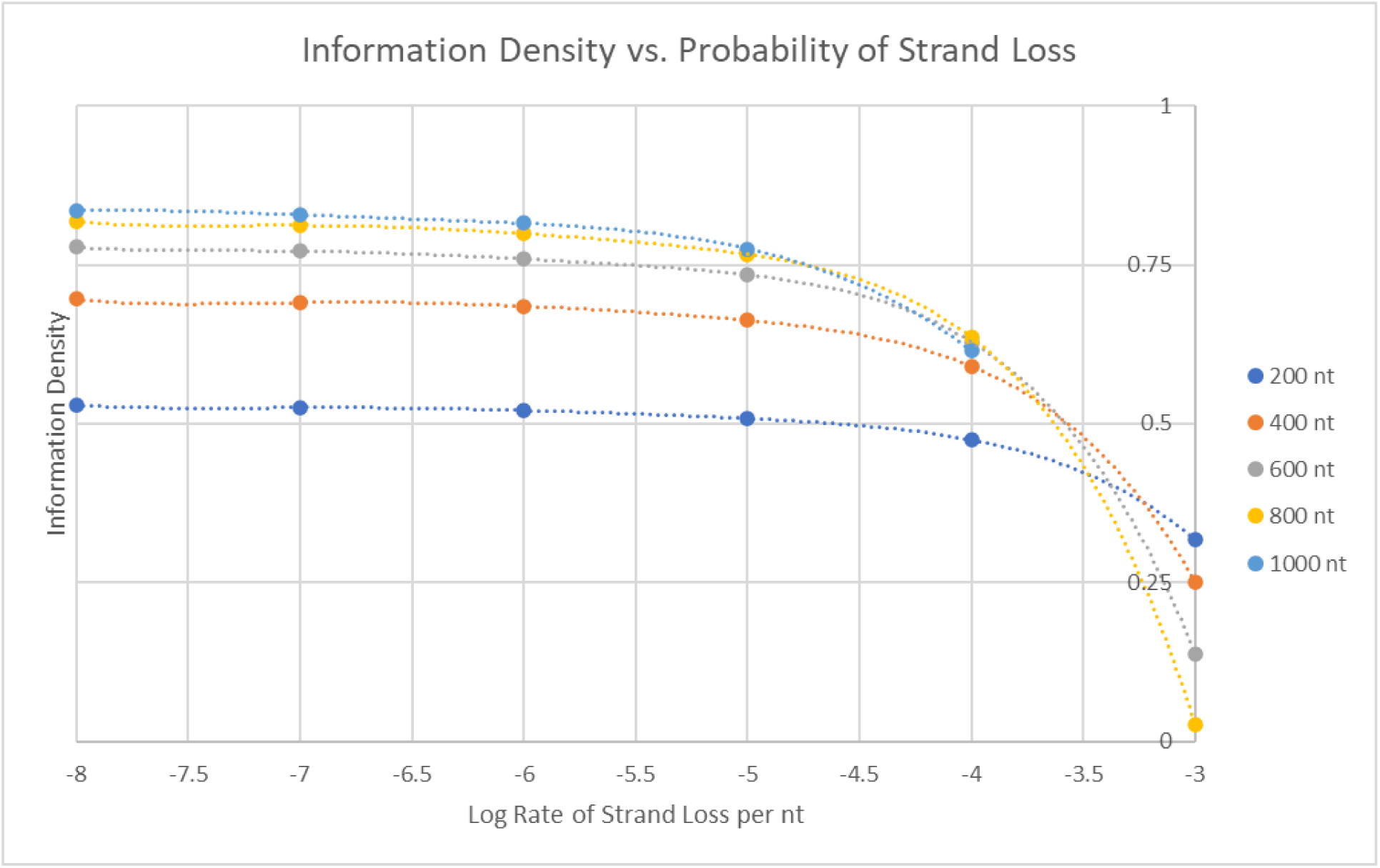
Correlation between the information density and the strand loss probability caused by breakage, in relation to strand length.

## Reed-Solomon Encoding

Reed-Solomon (RS) codes are frequently utilized in DNA storage systems due to their flexibility and robustness in dealing with errors.^47^ In this study, we employ RS codes to analyze the collective impact of strand breakage, strand length, and strand errors on both the overall stability and the storage density of DNA storage systems. While other codes exist, we utilize RS codes as an example to highlight general trends that should be taken into account when developing DNA storage systems. Though the intricacies of RS codes are beyond the scope of this paper, in essence, implementing RS codes gives storage systems the ability to detect and differentiate between two forms of errors: strand erasures (strand breakage or strand loss) with a probability of *p_strand erasure_* and strand error (mutations and deletions) with a probability of *p_strand error_*.^48^ RS codes incorporate a combination of both inner and outer codes. The inner code is designed to protect the DNA strand from internally occurring strand errors, while the outer code is intended to correct for strand erasures such as lost or defective strands (Figure 2). The code also has the ability to consider the existence of multiple copies of the same DNA strand, which means that a strand will only be considered lost if all copies of it are missing.^49^ This feature provides the flexibility to adjust the redundancy levels in the encoding scheme to attain a particular desired level of storage stability and error probability.

## Decoding Error Analysis

To determine a correlation between DNA system configurations and DNA stability, we examine the effects of strand loss caused by hydrolysis or mechanical breakage - a prevalent form of DNA degradation - on the probability of decoding errors, which occur when the original data can no longer be recovered due to errors or missing information. Specifically, we derived a mathematical model similar to Grass et al.’s to examine how the probability of decoding errors for an outer RS code is affected by strand loss. This was done by first assuming a linear relationship between strand length and the likelihood of strand breakage based on previous experimental data.^51^ Then, we make the assumption that the probability of strand loss can be exponentially reduced by having multiple copies of the same strand. This is because for an erasure to occur, a specific strand must lose all of its copies. Therefore, we approximate the relationship as follows:

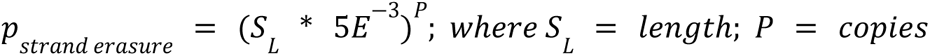

This formula was derived in a way that ensures DNA strands up to 1000 nucleotides (nt) in length have an erasure probability of 50% or lower, but this threshold will need to be adjusted to reflect the stability of the DNA storage system being examined. For our purposes, we set the error probability to 1.0*E*^−2^ and 1.0*E*^−3^ per nucleotide, which covers typical rates of synthesis and sequencing errors.^52 53^

Figure 3 illustrates the effect of increasing strand length on decoding error probability across different storage configurations, including variations in the number of strand copies and *p_strand error_* rates. The x-axis represents the strand length while the y-axis displays the logarithmic decoding error probability. To facilitate comparison, we employ a system with a read error rate that resembles that of hard drives, estimated to be around 0.1*E*^−14^/*bit*, with an error probability of – 1.5*E*^1^.^54^ Our analysis indicates that strand length is a critical factor in the likelihood of strand breakage. Longer strands have a higher probability of experiencing strand loss, leading to a significant increase in the probability of decoding errors. Other factors such as lower *p_strand error_* values can notably decrease the probability of errors, while higher strand copy numbers significantly lowers the probability of losing every single copy of a specific strand. Although experimental systems currently require large copy numbers, our findings suggest that copy number requirements for future systems can be lower than expected, especially if strand lengths are limited within the range of 200 to 400 nt.

## Information Density Analysis

In order to investigate how altering information density, strand loss, and strand length affect one another we lowered the strand breakage probability from 1.0*E*^−3^ per nucleotide to 1.0*E*^−8^ and then calculated the information density for various strand lengths. Our results are presented in Figure 4, where the strand loss probability is displayed on the x-axis, and the information density on the y-axis.

According to our results, the system exhibits higher information density with shorter strands when the probability of strand breakage is high. This is because shorter strands are less prone to breakage, and as a result, they require fewer error correction encodings, leading to higher information density. Longer strands, on the other hand, are more likely to experience breakage, resulting in a need for more error correction, which ultimately lowers information density. However, as the rate of strand breakage lessens, longer strand lengths become advantageous since more information can be stored within every strand, and the RS code will still be able to operate efficiently even with fewer error correction encodings. After conducting further analysis, we discovered that the correlation between ideal strand length and information density can differ significantly depending on the extent of strand breakage (Figure 5). For high rates of strand loss, shorter strands are not always the best option. Instead, there may be an ideal strand length that strikes a balance between the required overheads in each sequence (file indices and addresses) and the optional encoding overhead used for strand loss compensation.

**Figure 5.**
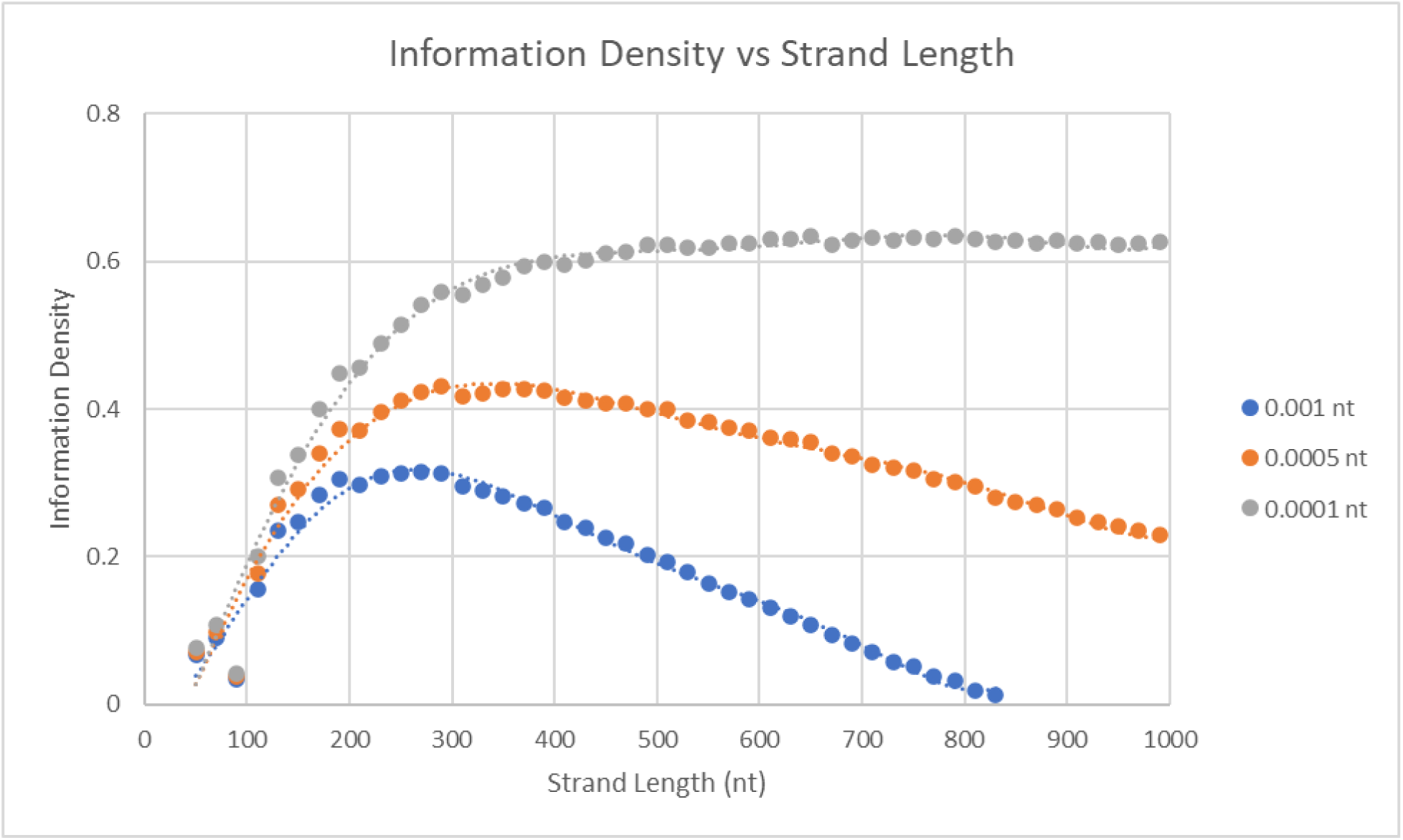
Correlation between the DNA storage system’s information density and strand length configuration, in relation to the strand breakage probability.

The results of this study highlight the importance of tailoring system parameters, such as copy numbers and strand length, to specific scenarios, including those where error rates may be affected by environmental factors and the intended use case of the storage system. For instance, for an Archival Storage system whose DNA is enclosed in silica, the probability of strand loss or breakage is much lower, thereby enabling the use of longer DNA strands and higher information densities. Conversely, for Working or Short-Term Storage systems, shorter strand lengths and lower information density requirements would be more appropriate due to the higher likelihood of strand loss.

## Future Prospects

Ensuring that data retrieval is successful and dependable is a crucial requirement for information storage systems.^55 56 57 58^ While there is some understanding of DNA stability under various conditions, the measurements often exhibit considerable noise and variation among researchers, experimental groups, and even within individual experiments. Factors such as DNA handling and confounding variables like DNA solubility can impact measurement accuracy. To increase confidence in the effectiveness of DNA storage technologies, it would be beneficial to conduct experiments that consider a wider range of parameters. Fine-tuning these system configurations and factoring in sources of variability will be essential for developing reliable commercial DNA storage products.

In addition, the precision of current techniques used to assess DNA degradation, such as mass measurements and quantitative PCR, is limited. Various forms of DNA degradation can occur, including biased loss of DNA strands based on properties like base content, specific sequences, or length, and chemical alterations like depurination that cannot be directly measured by sequencing or QPCR-based methods. It is crucial to understand how environmental conditions affect each of these degradation mechanisms to design a reliable DNA storage system.

Improving the efficiency of DNA storage systems requires a better understanding of DNA stability. However, evaluating DNA stability is challenging as it can be measured using various methods, such as strand breakage rate, mutational rate, and strand loss. In addition, DNA degradation rates have been observed under varying conditions, such as temperature, environmental, temporal, and buffer conditions. Furthermore, the resulting degradation effects can also vary based on the storage system, including factors like the DNA strand length, storage density, and encoding techniques. Yet despite the limited knowledge of DNA stability, it is still possible to develop reliable DNA storage systems using existing technologies. However, it is essential to understand the limitations and tradeoffs in system configurations and employ models to support them. Improving the precision of measurements, developing novel technologies, understanding mechanisms of degradation, and refining encoding techniques will all contribute to enhancing the efficiency and reliability of DNA-based information storage, making it more viable for commercial use.

## APPENDIX 1: Stability Simulation Code

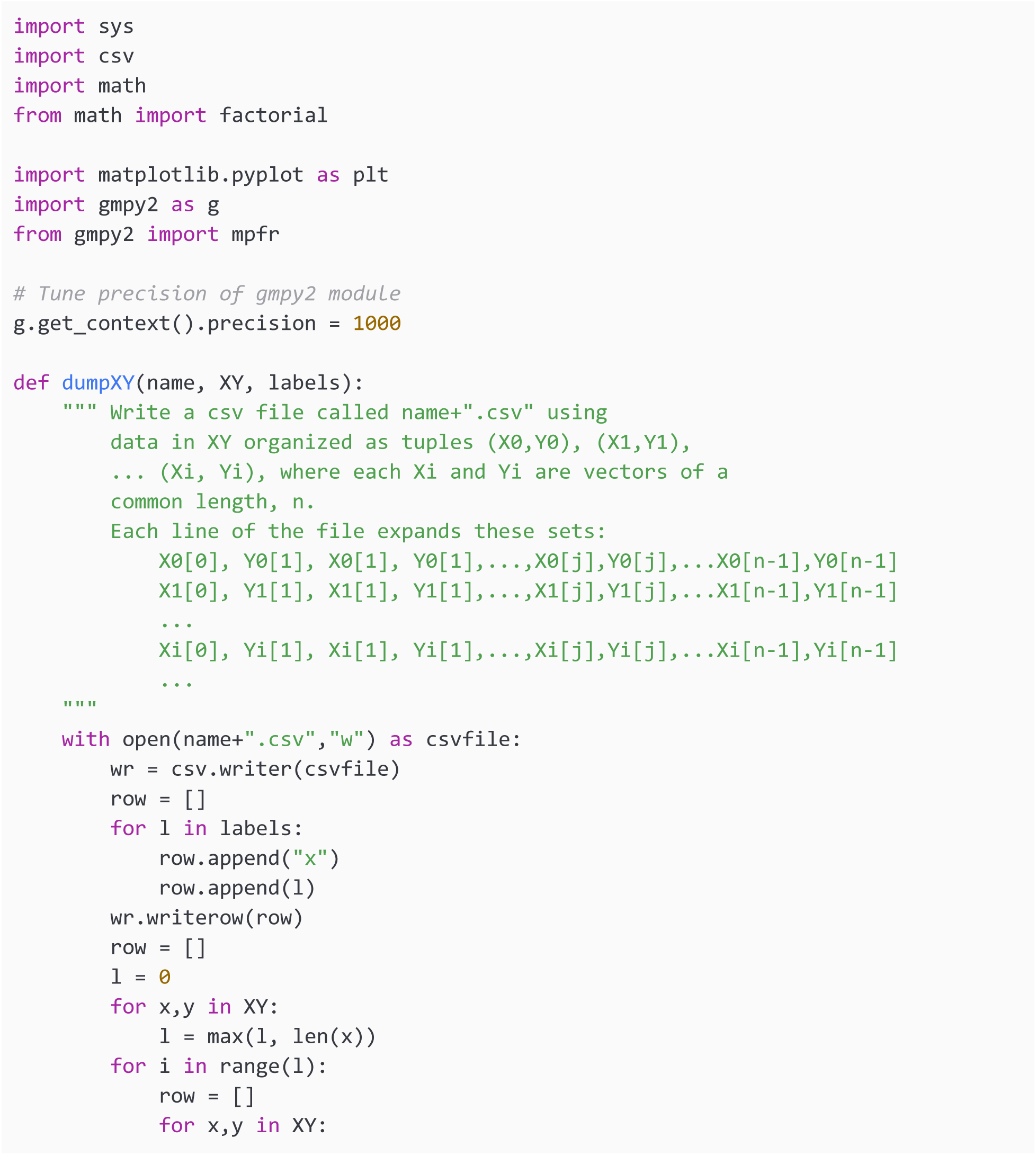
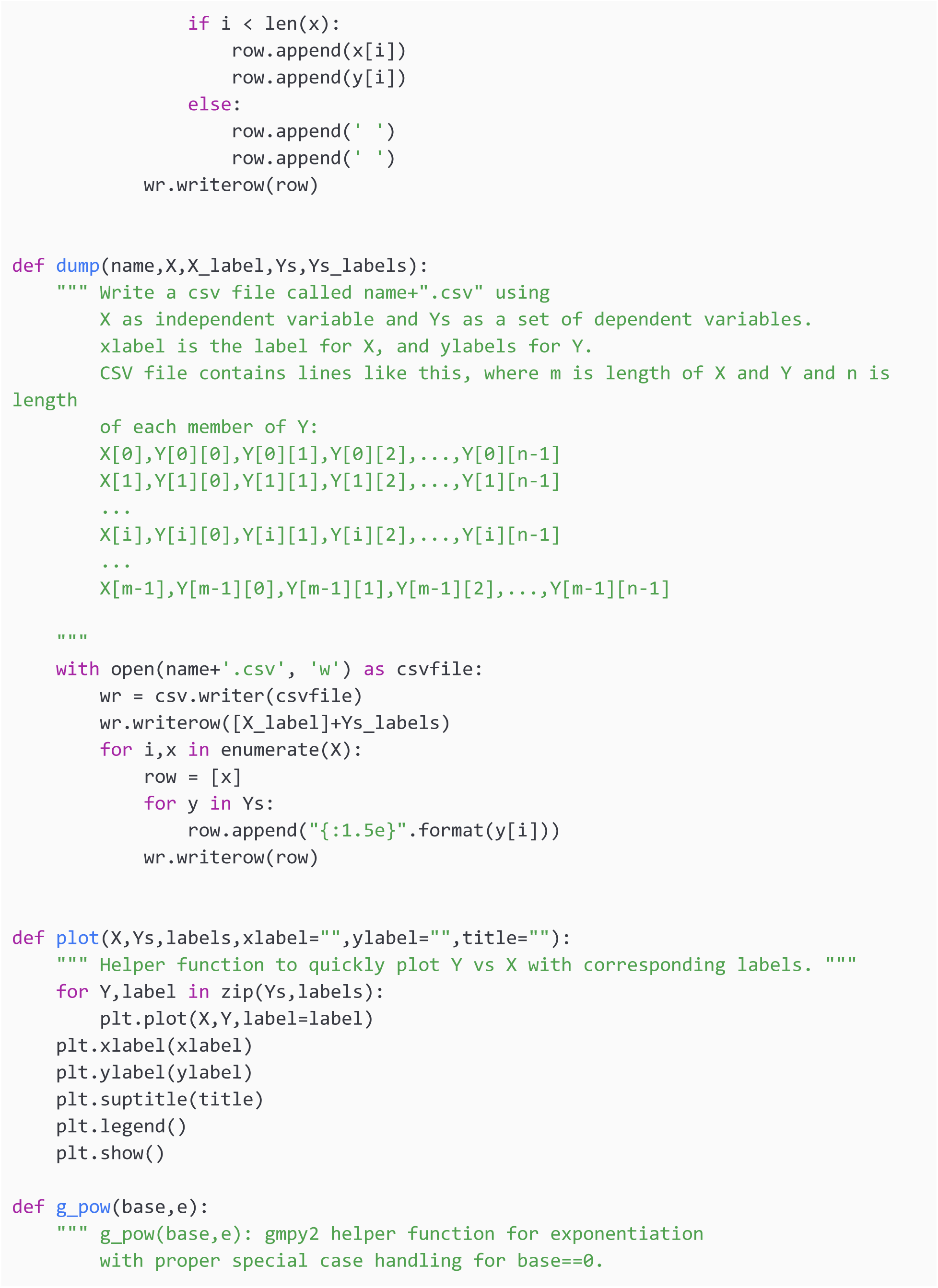
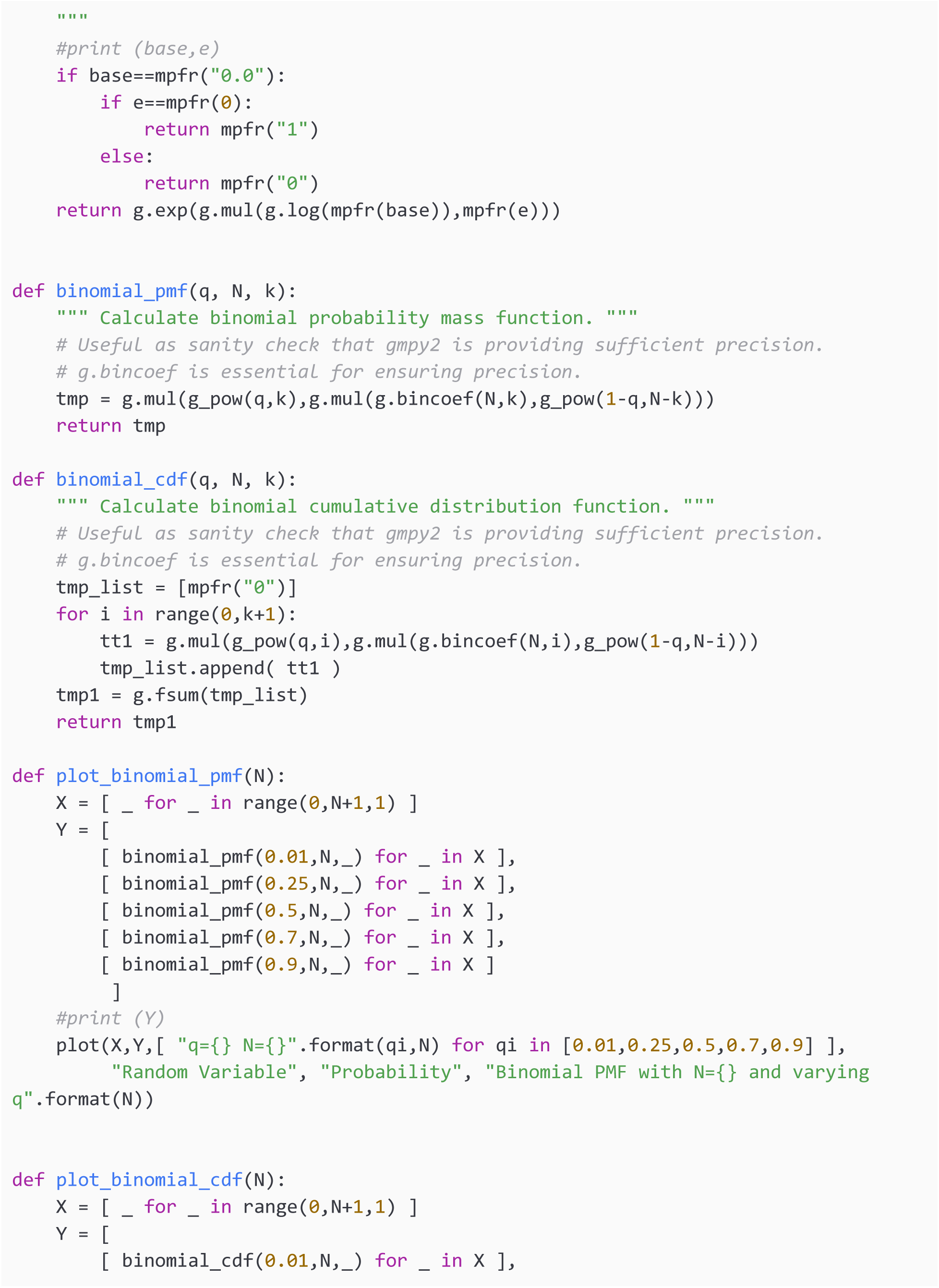
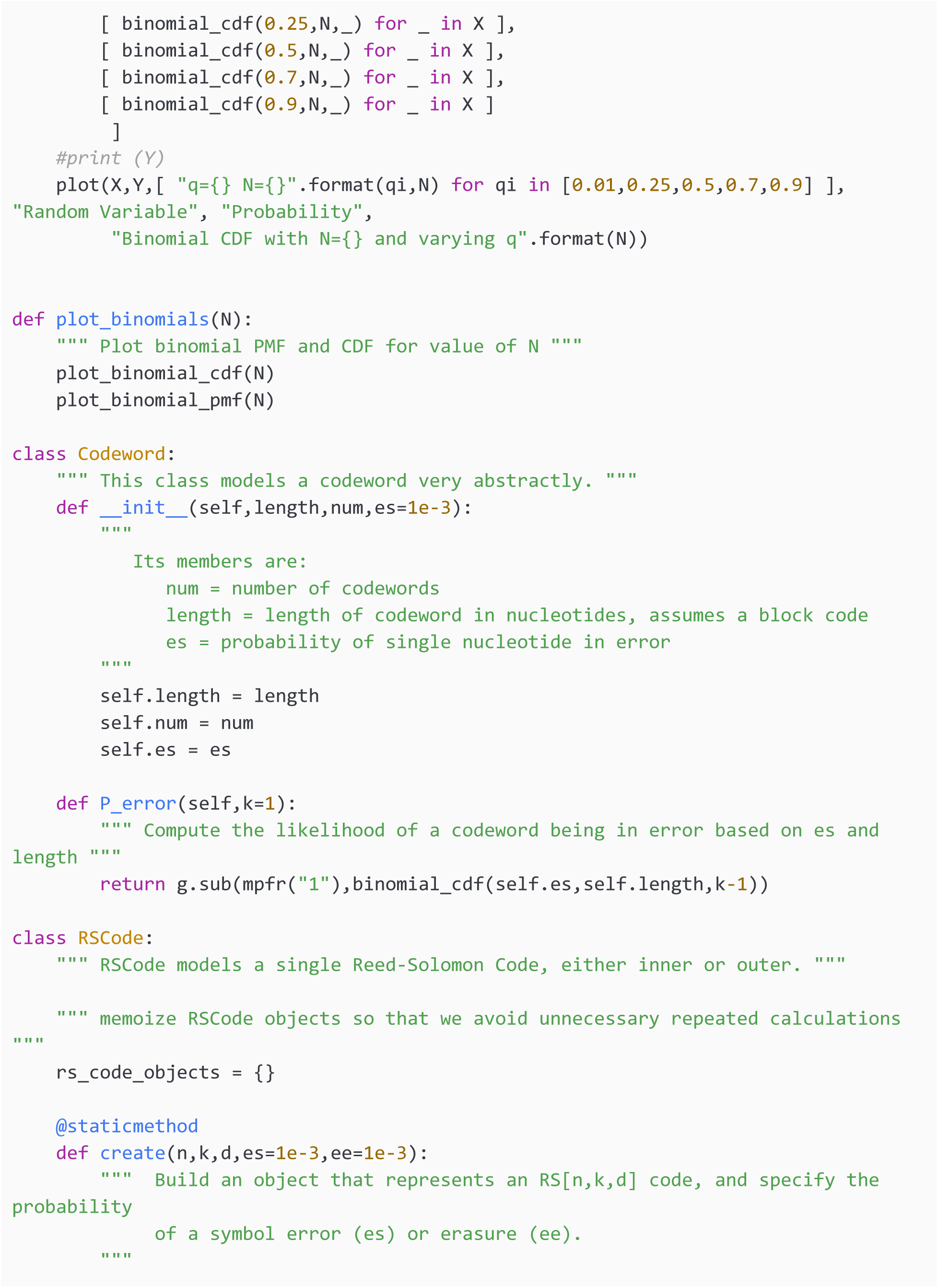
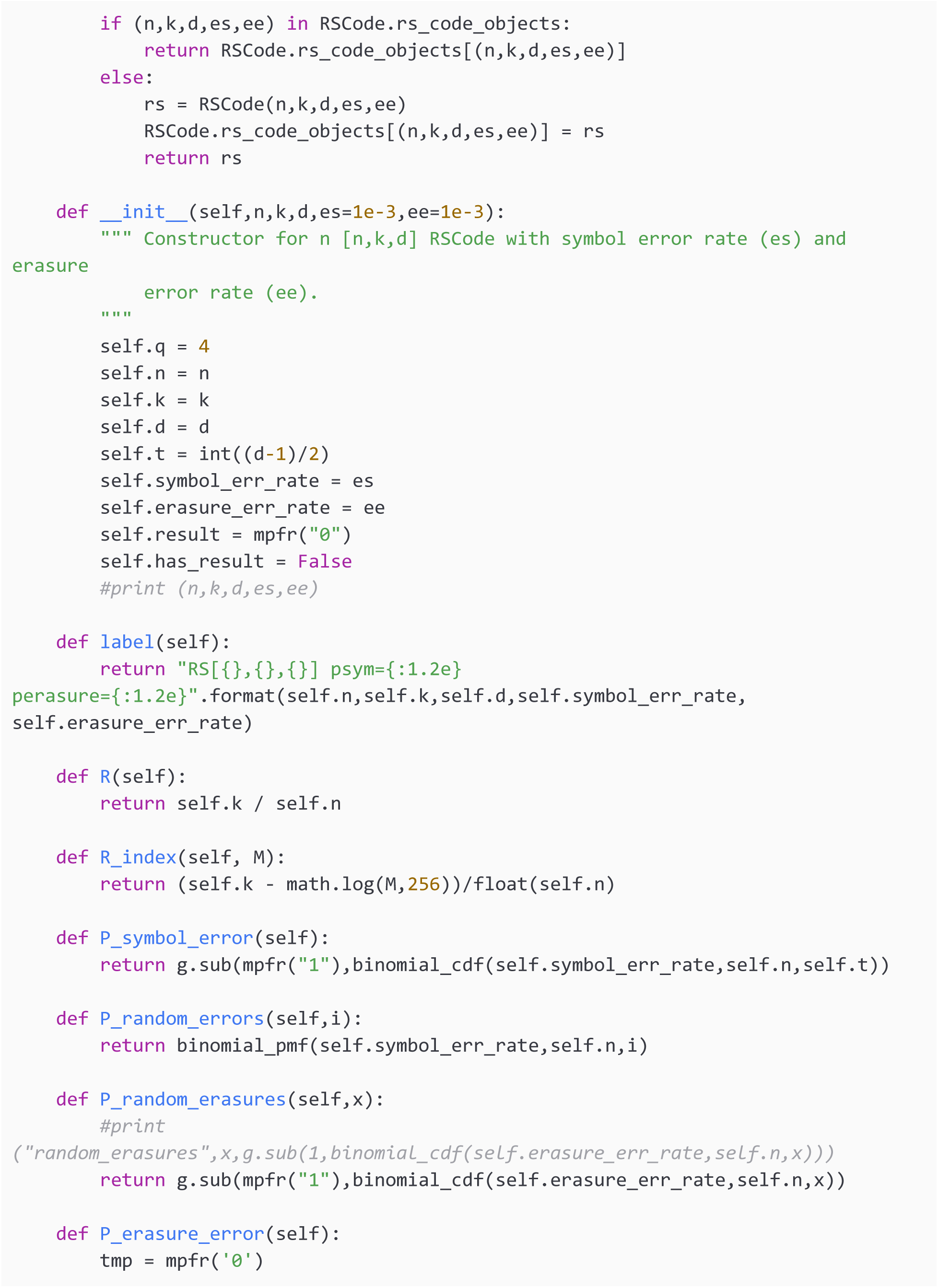
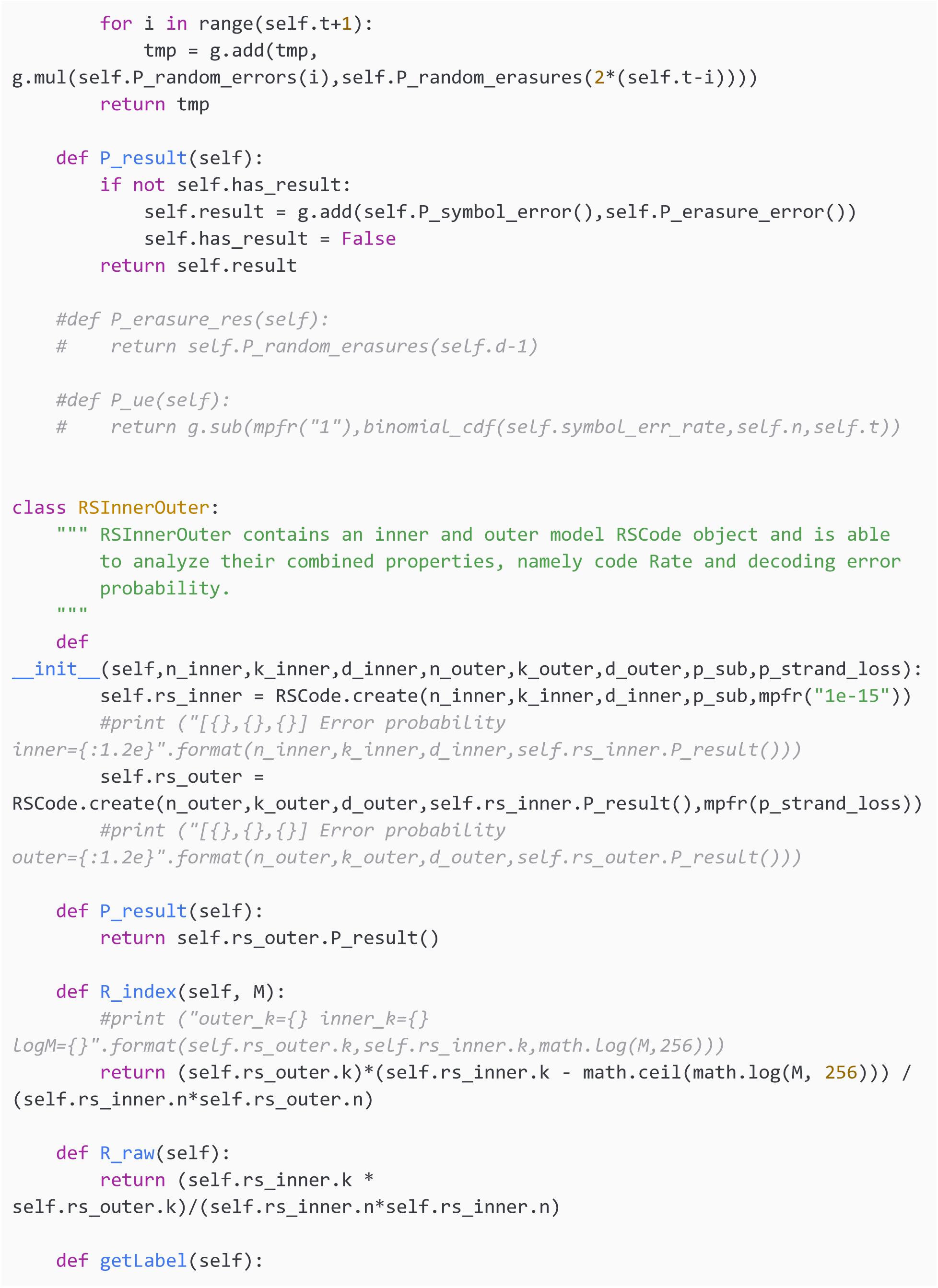
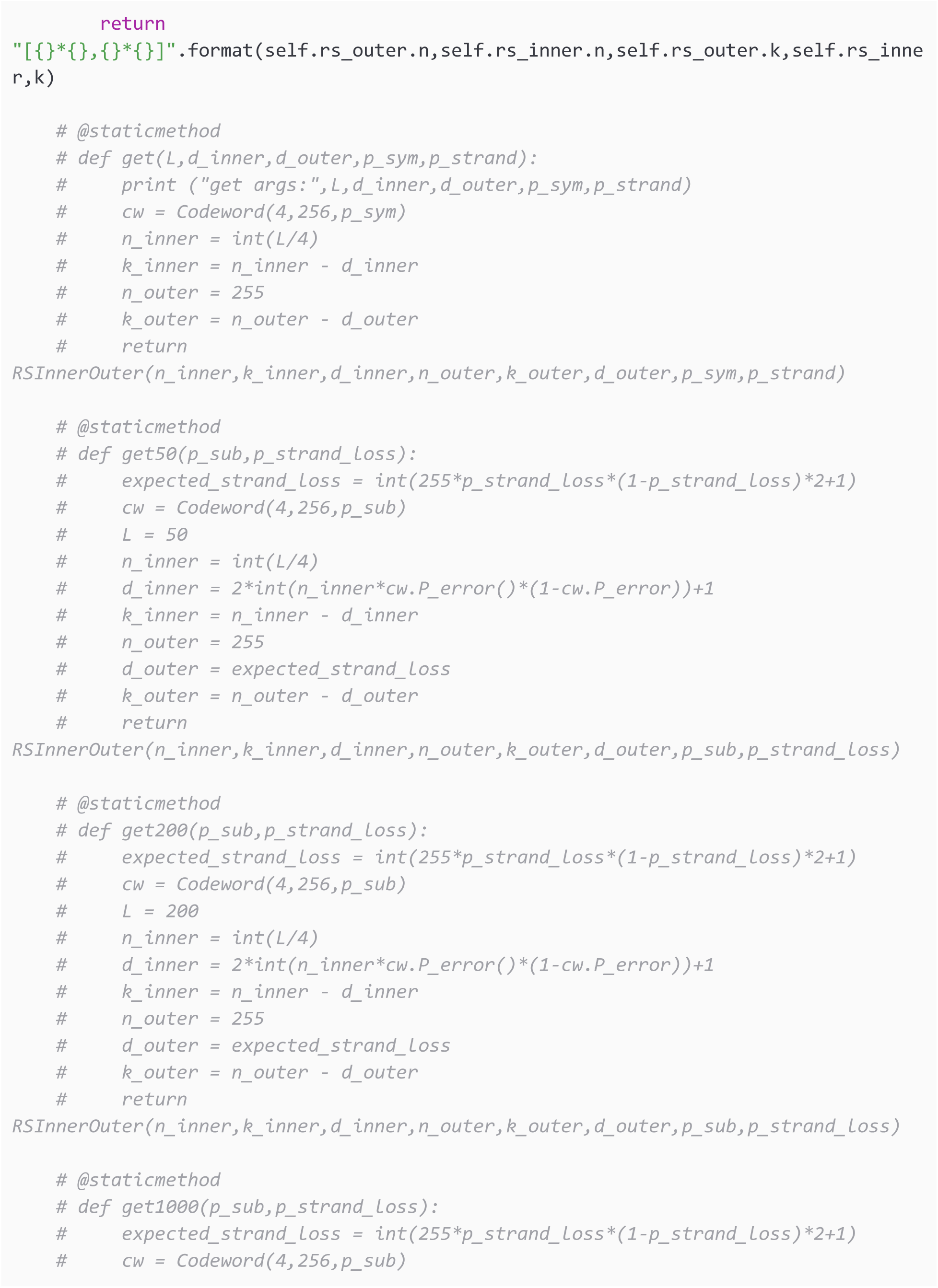
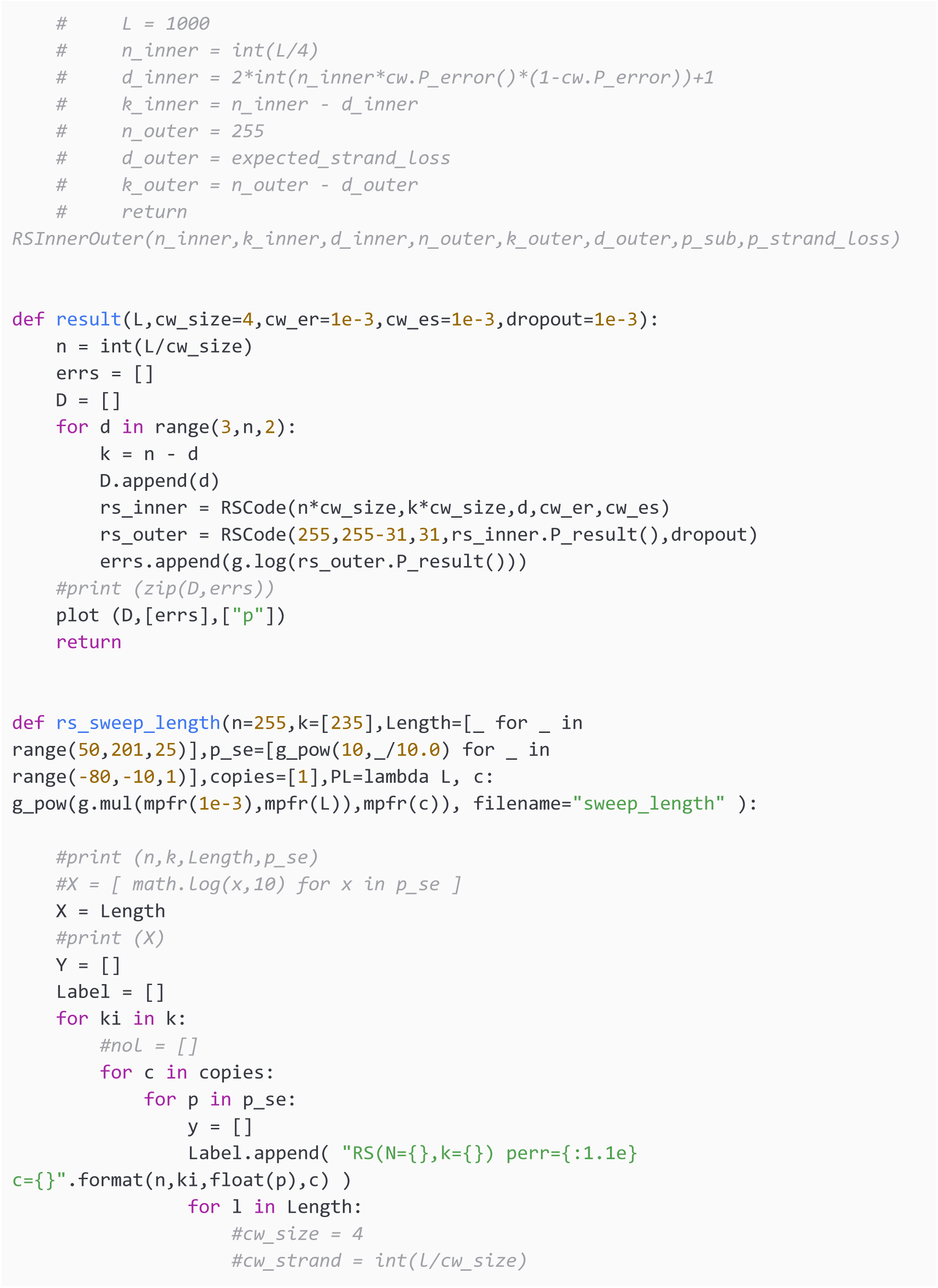
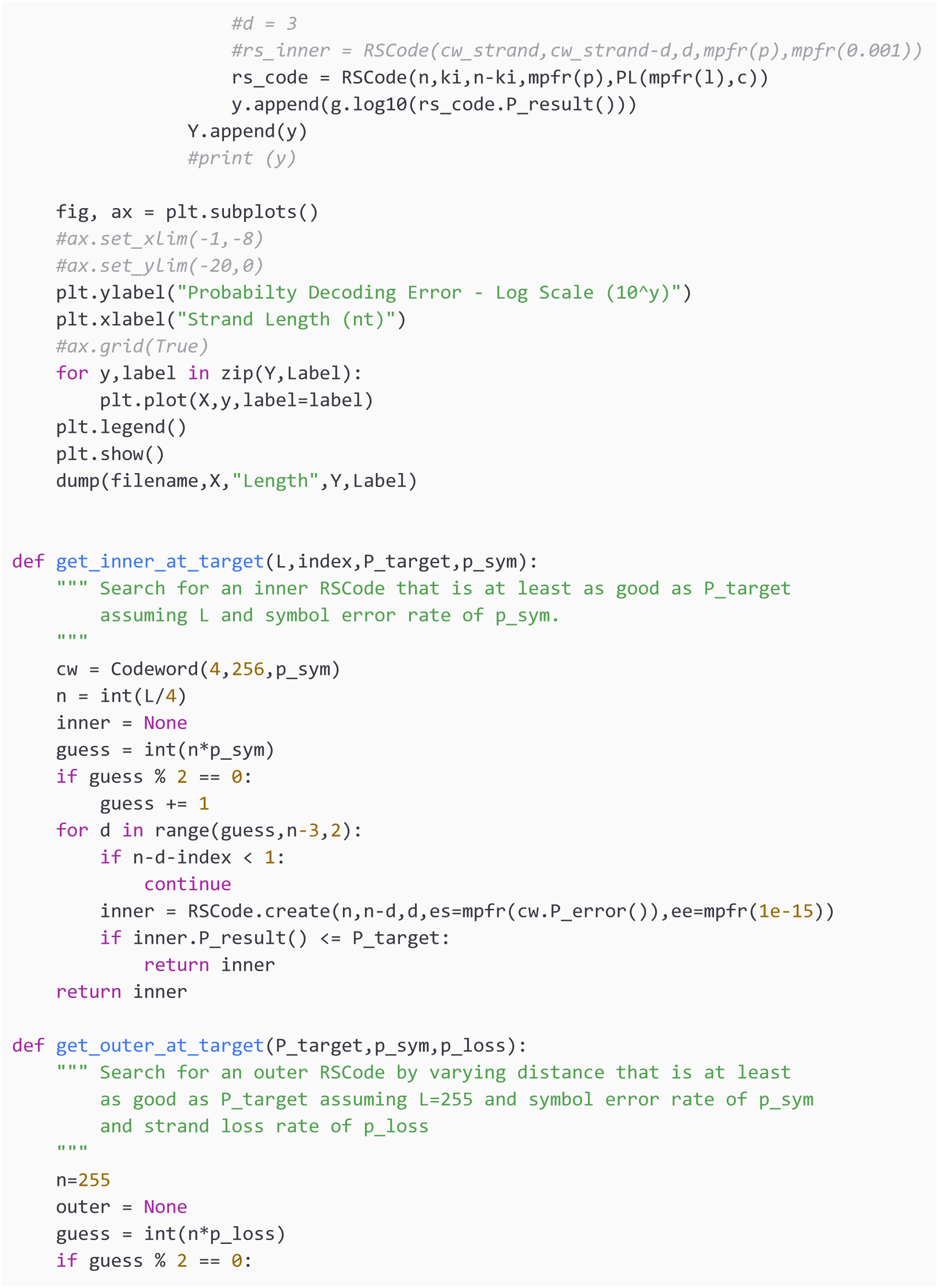
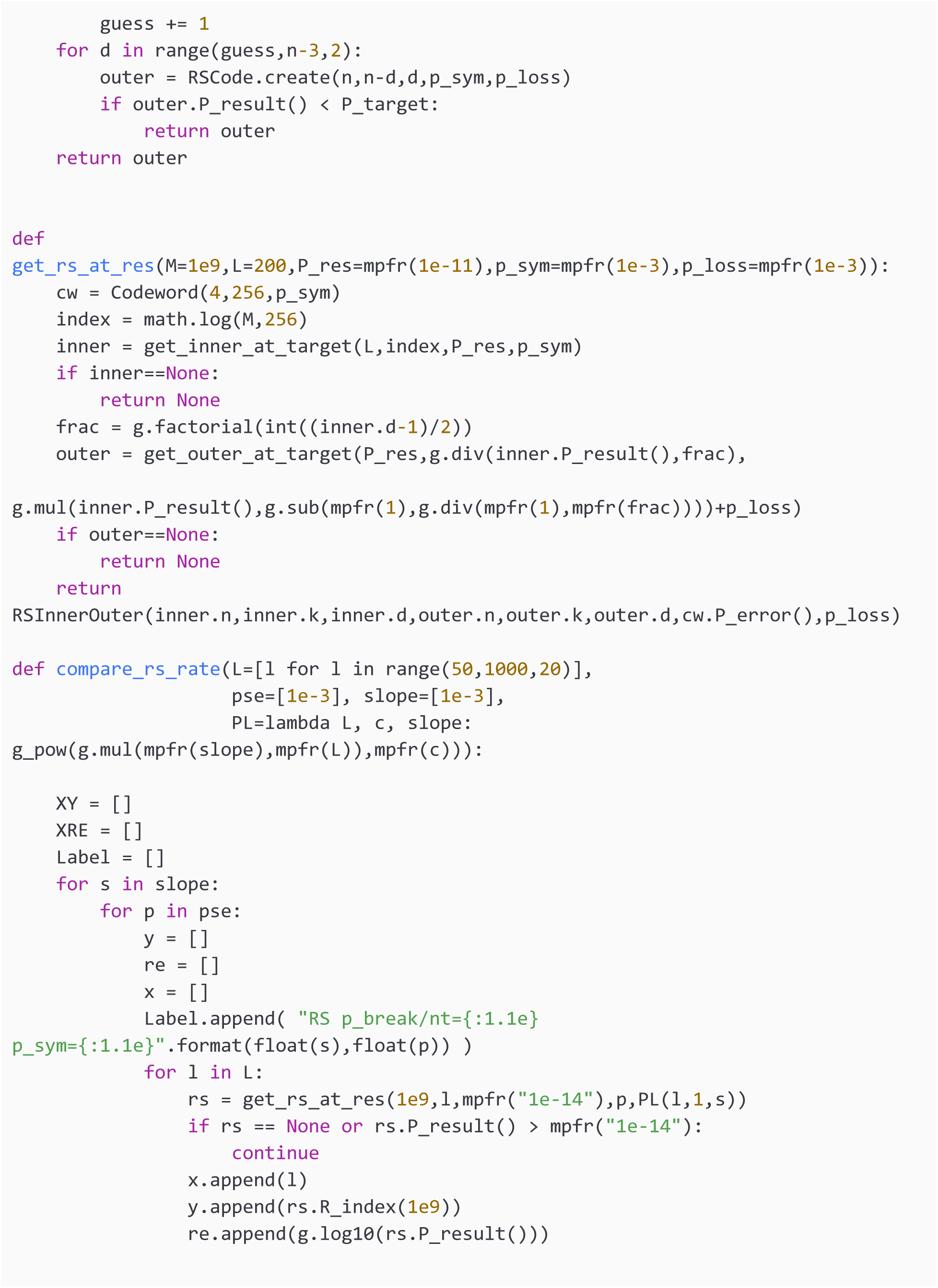
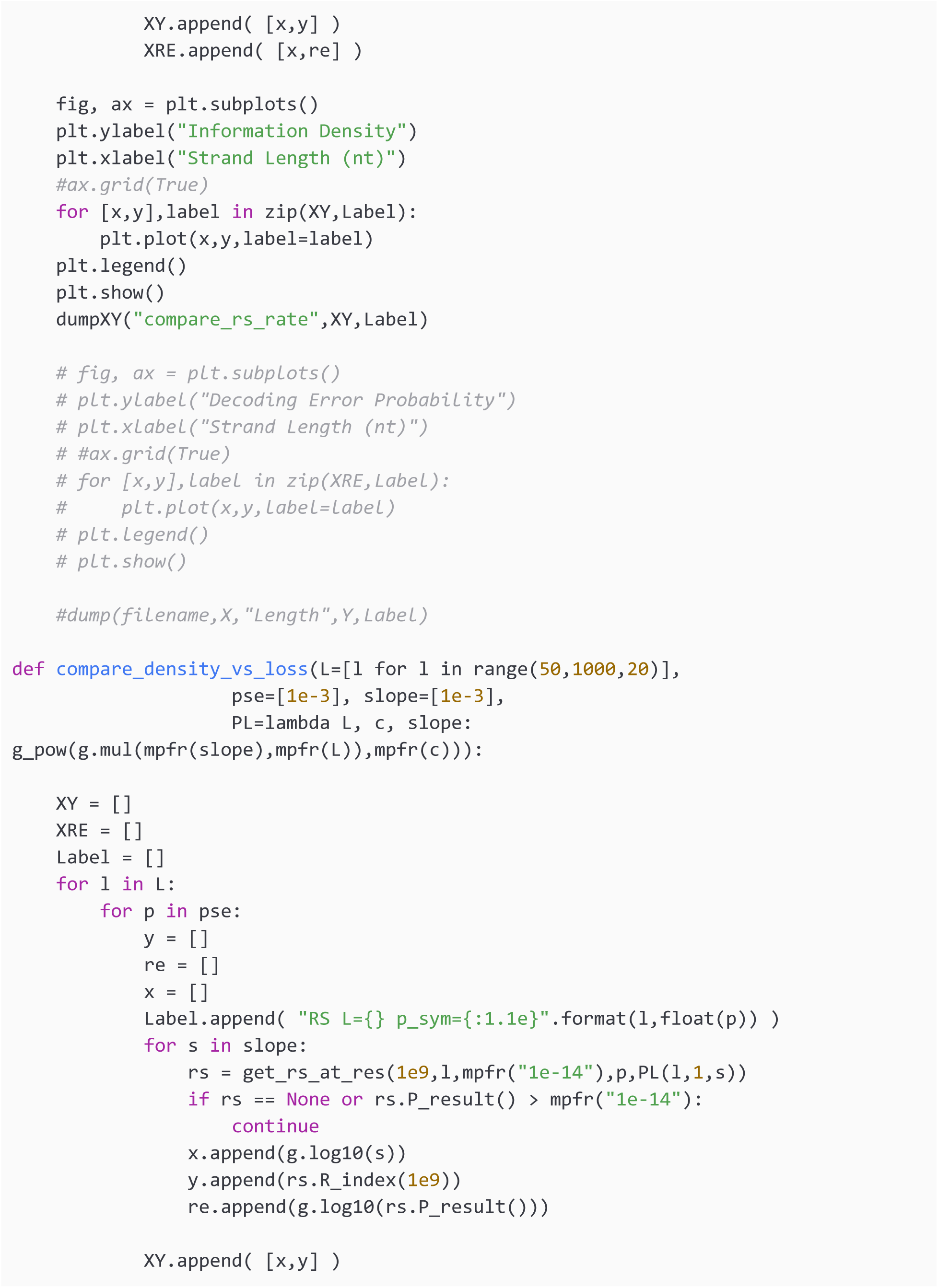
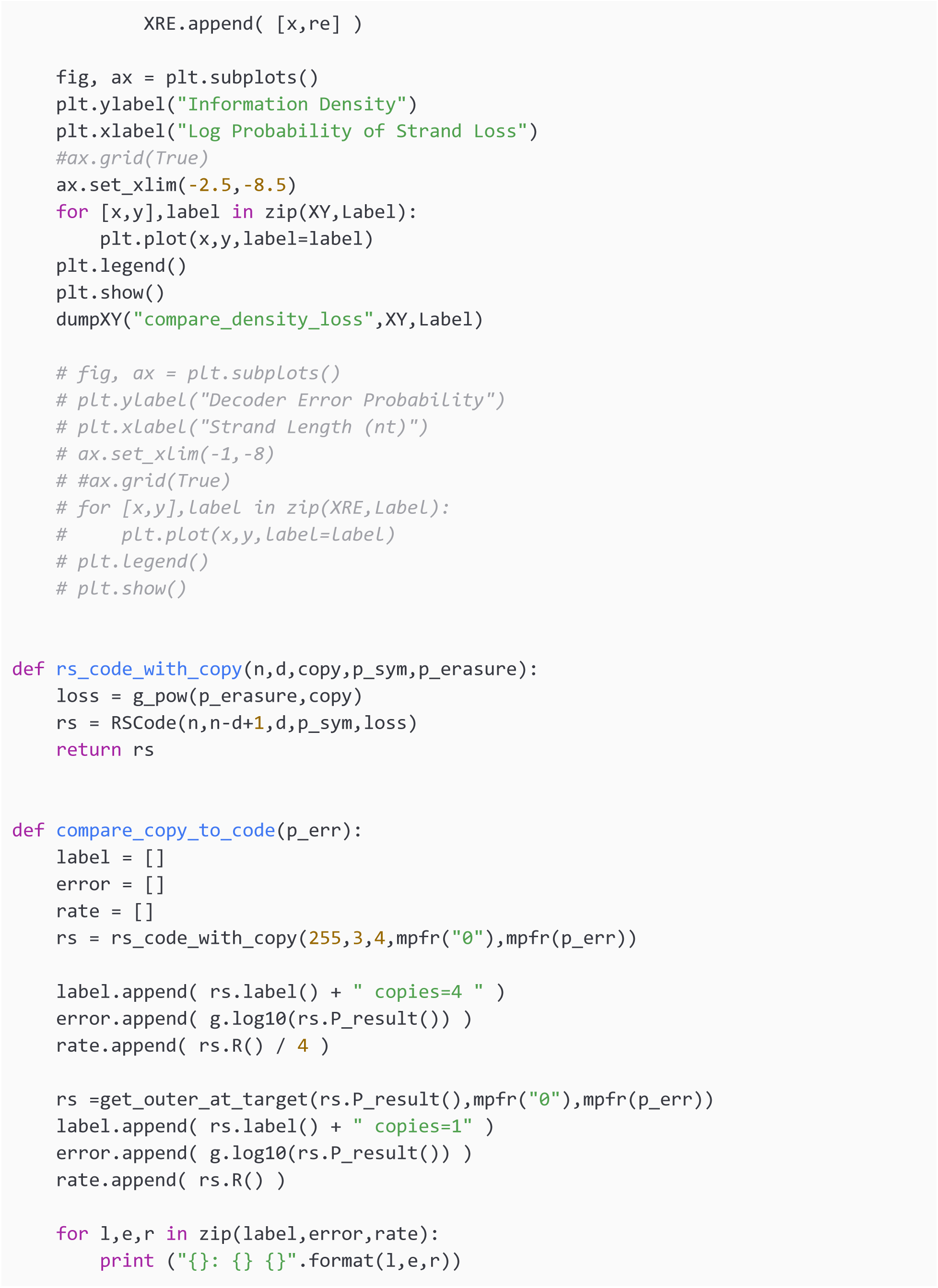
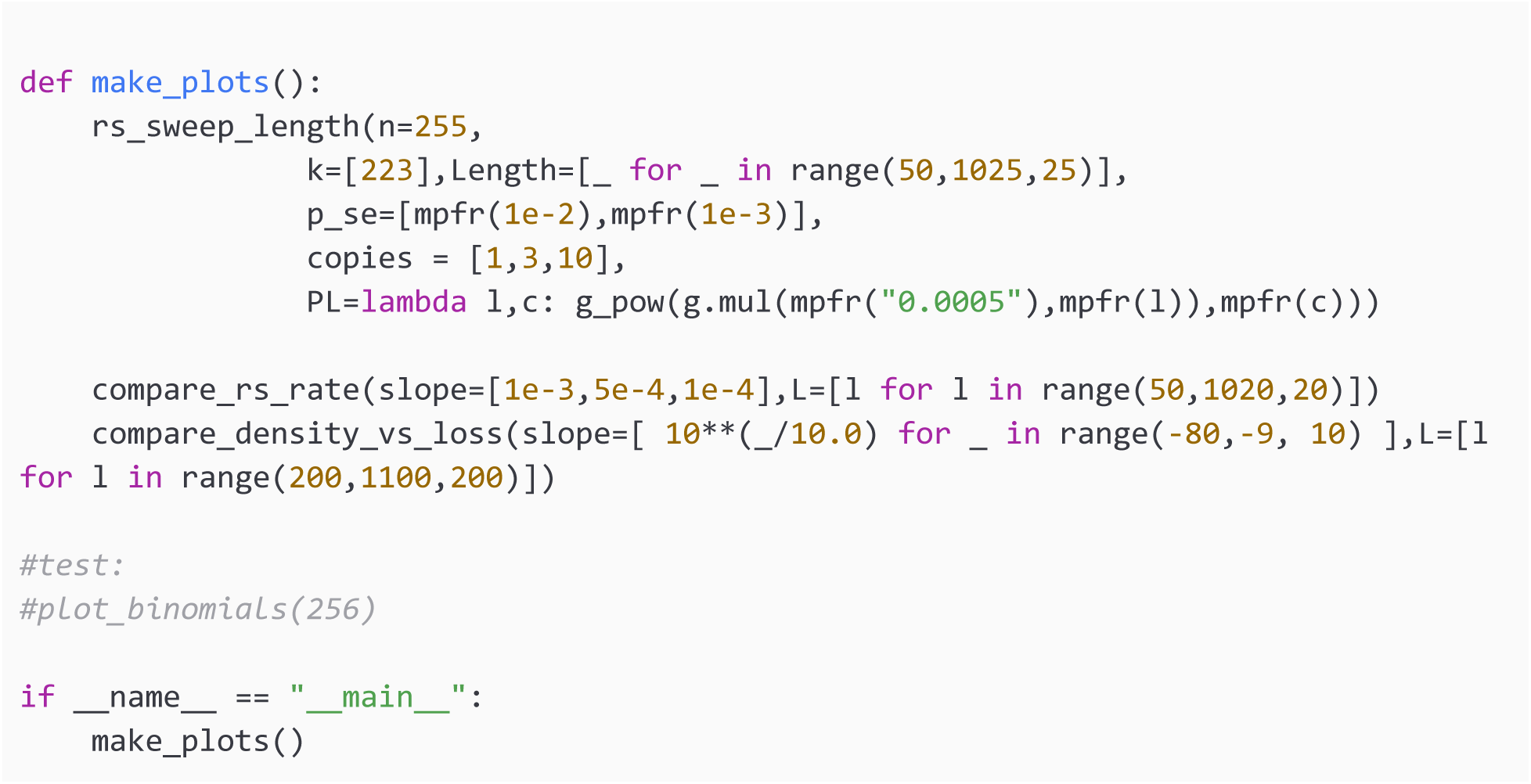

## APPENDIX 2: Raw Dataset Availability

### 2.1 Information Density Loss

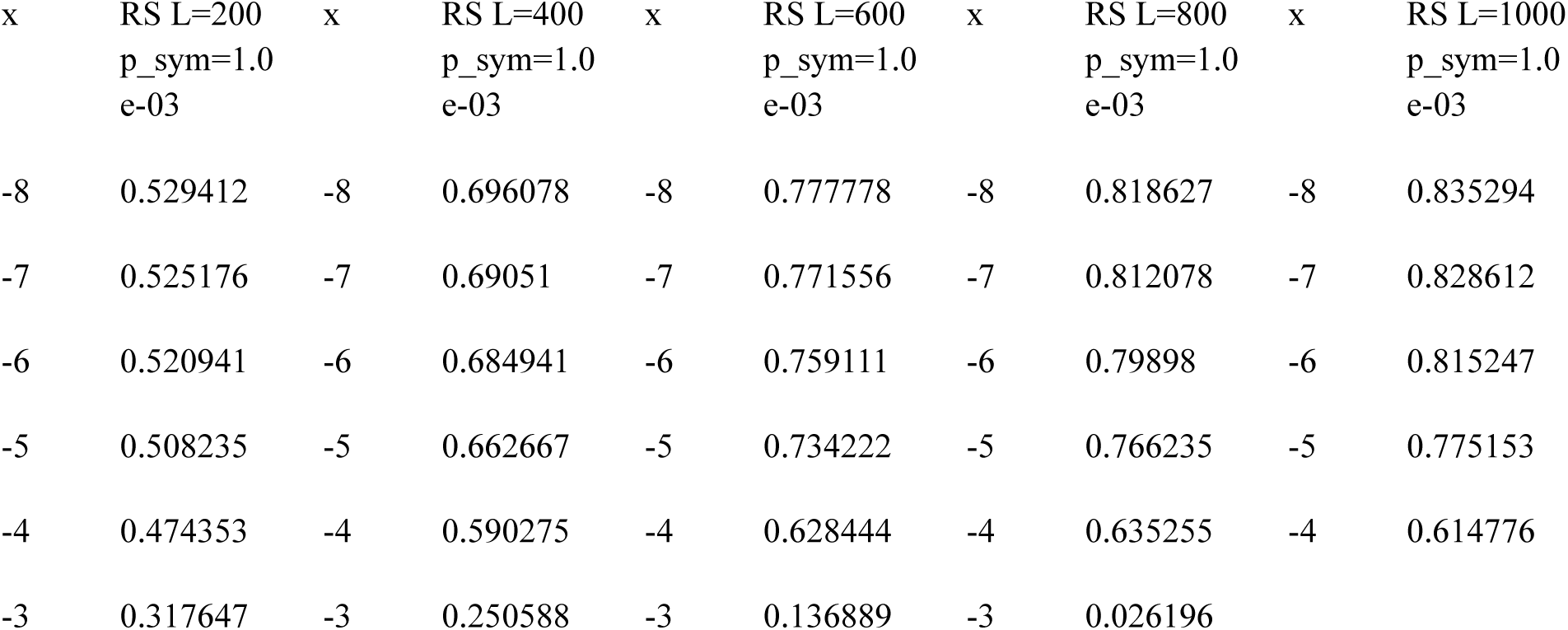

### 2.2 RS Strand Breakge Rates

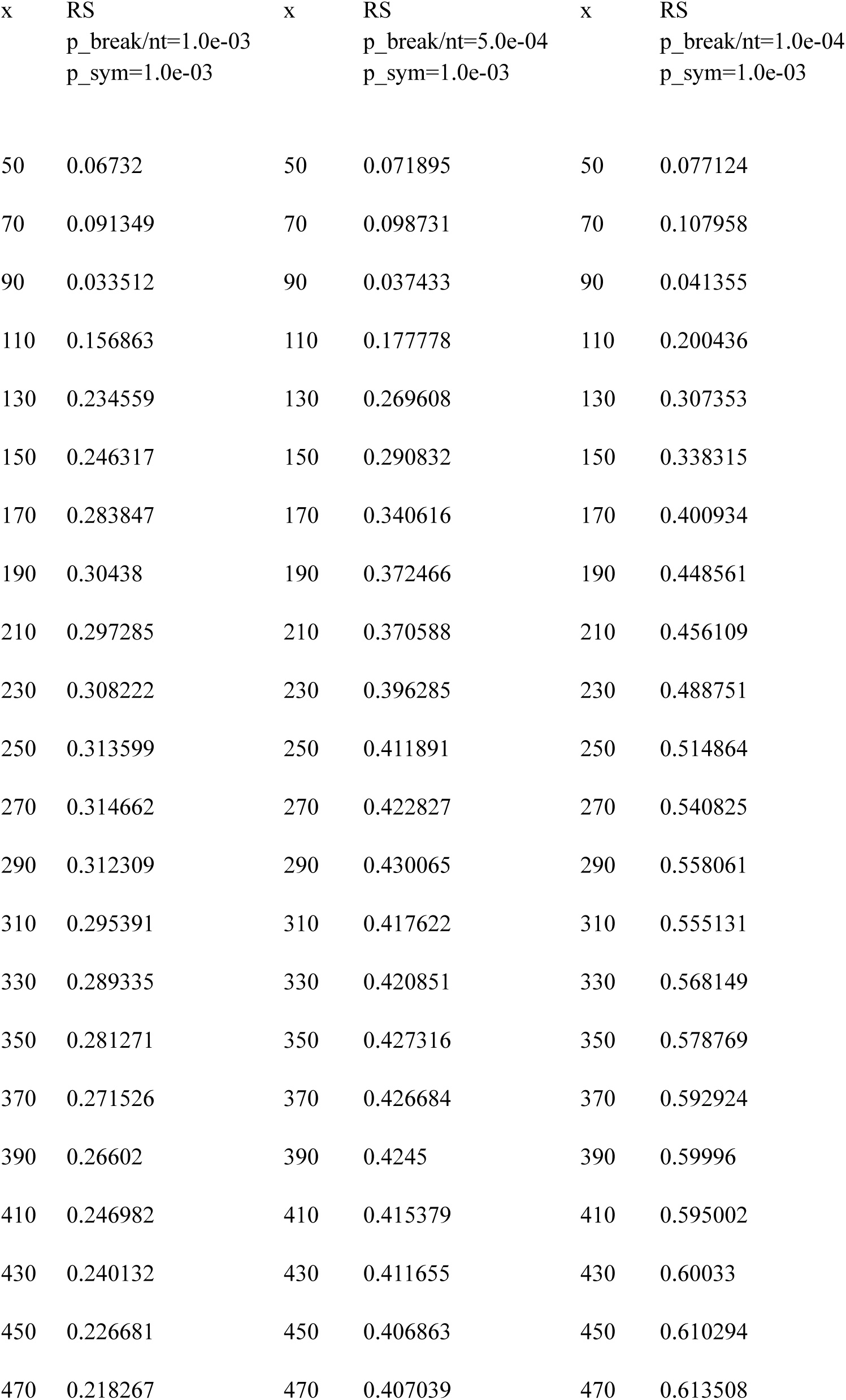
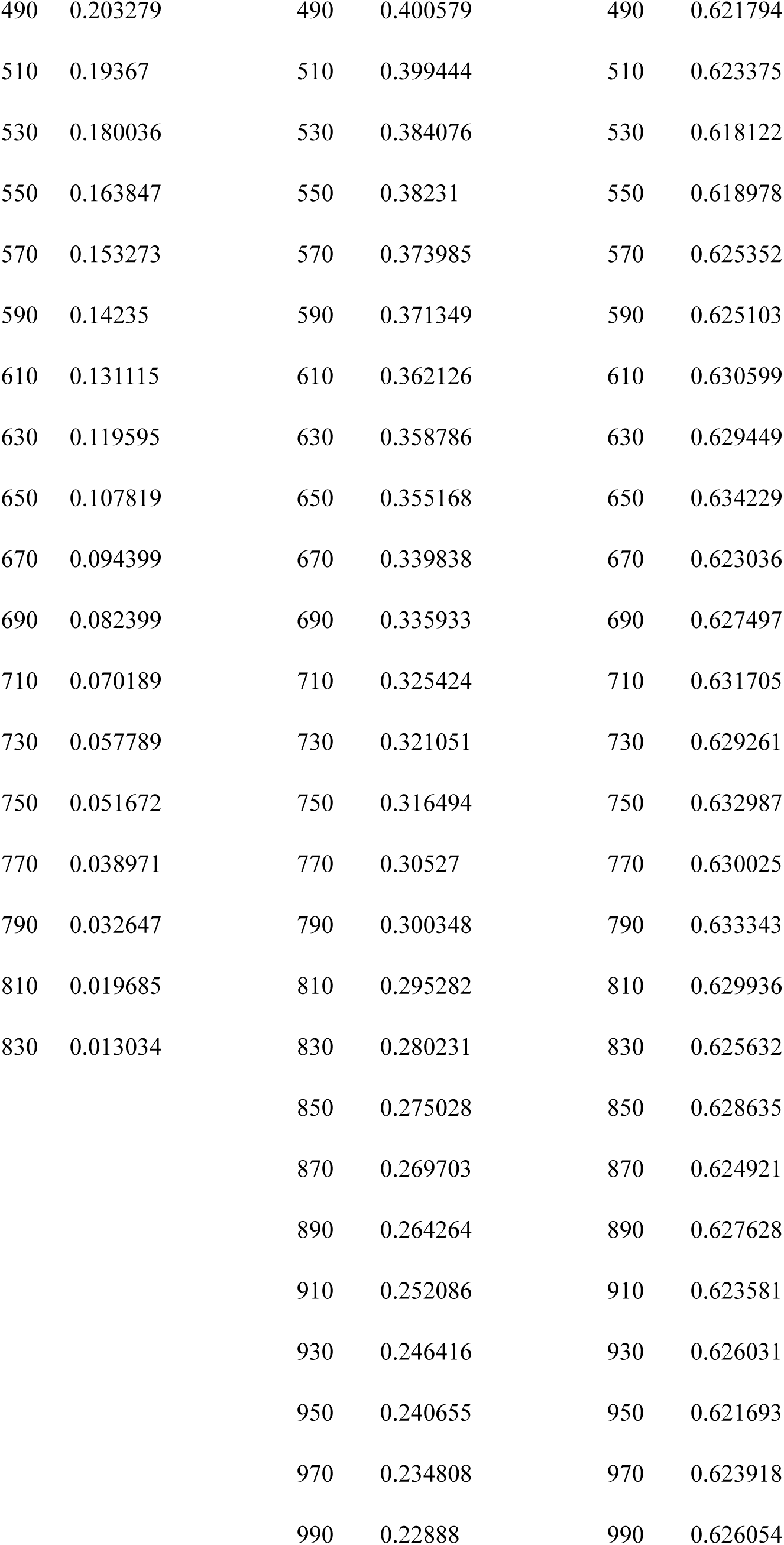
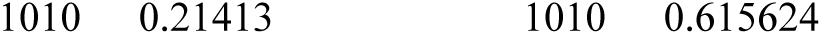

### 2.3 Strand Length Configurations

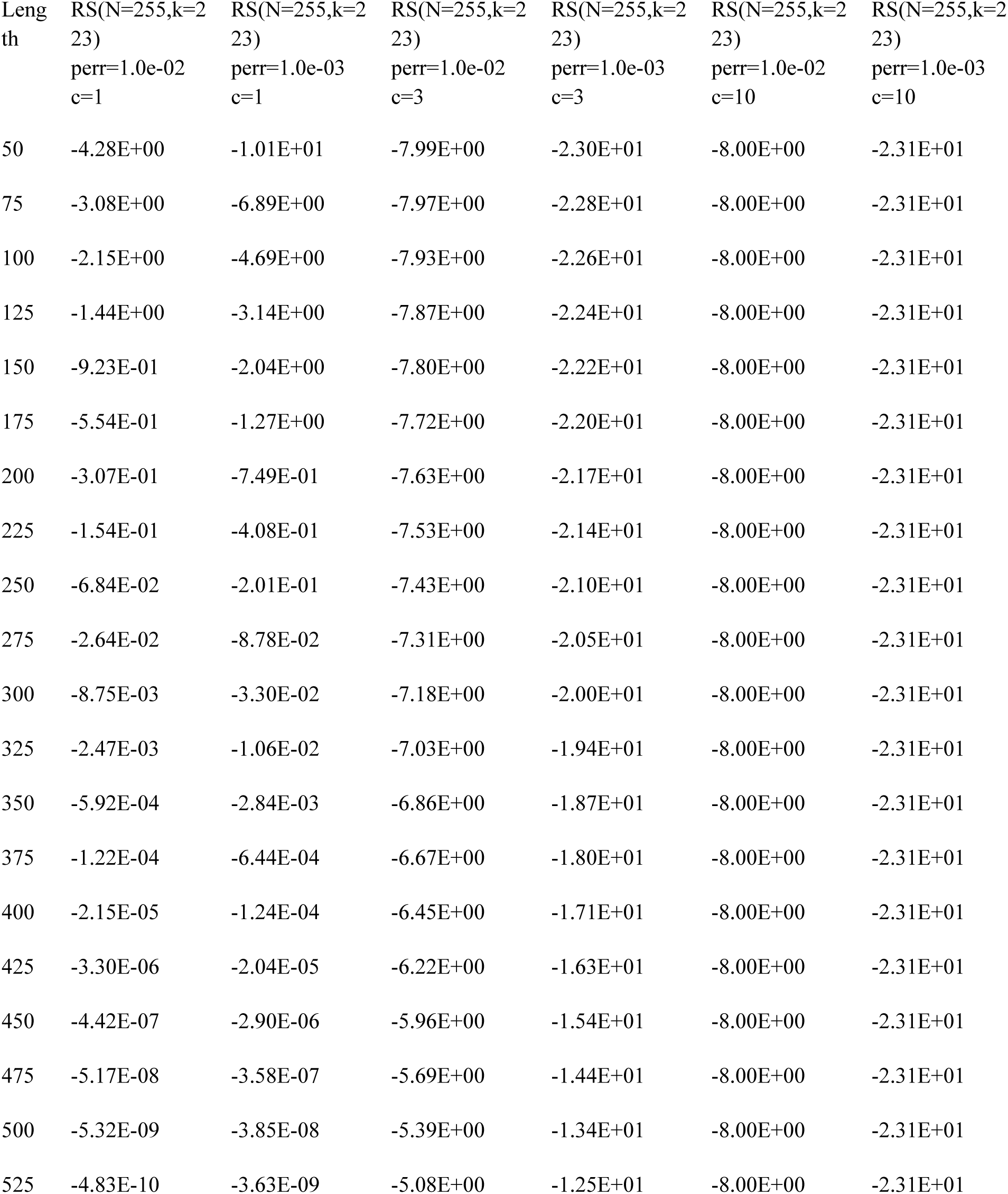
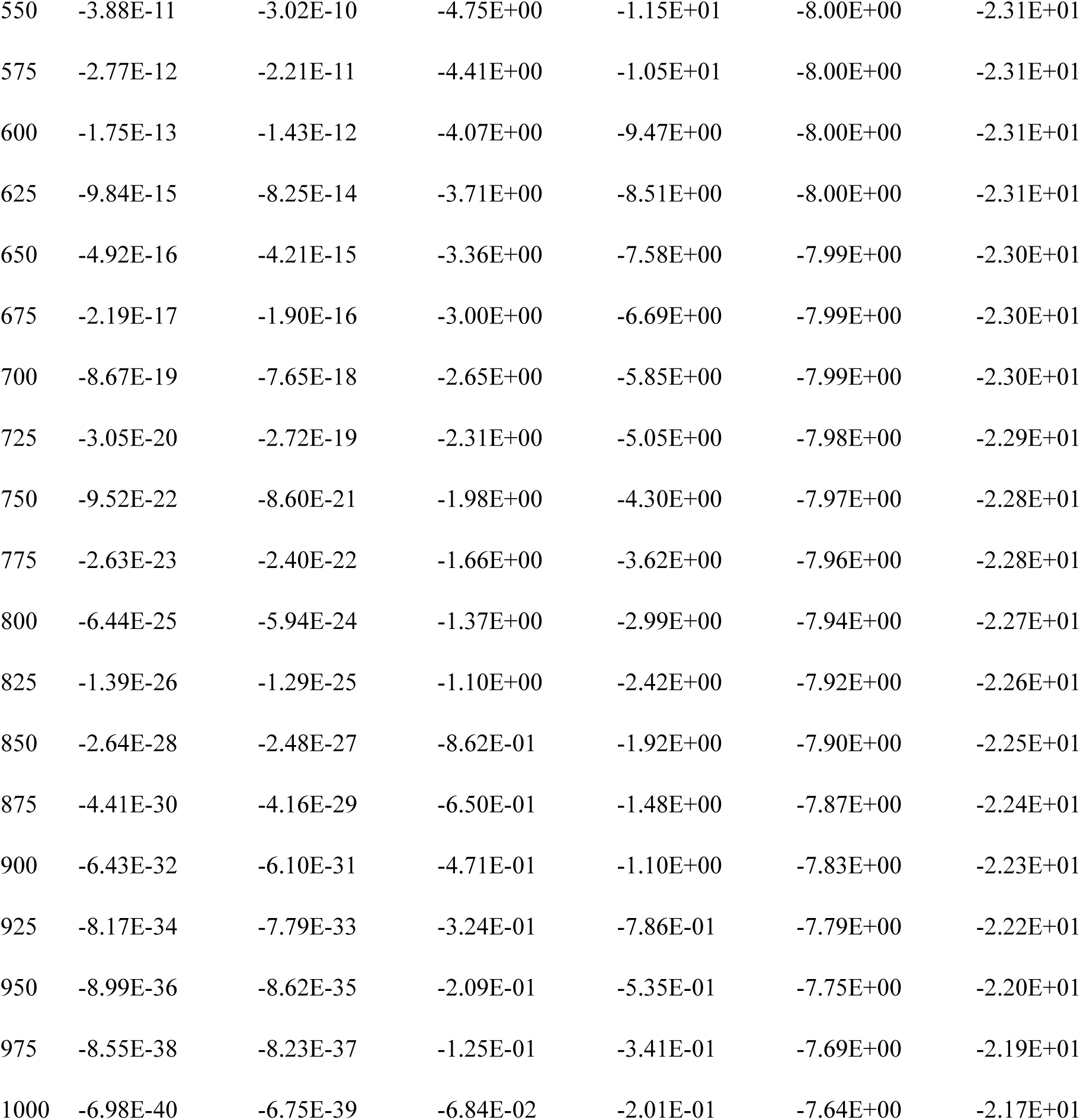

## Footnotes

1 Cao B, Zhang X, Cui S, Zhang Q. Adaptive coding for DNA storage with high storage density and low coverage. *npj Syst Biol Appl*. 2022;8(1):1-12. doi:<u>10.1038/s41540-022-00233-w</u>

2 Yupu Z, Myers D. Zettabyte reliability with flexible end-to-end data integrity | IEEE Conference Publication | IEEE Xplore. IEEE Xplore. Published May 2013. Accessed March 21, 2023. https://ieeexplore.ieee.org/document/6558423

3 Koot M, Wijnhoven F. Usage impact on data center electricity needs: A system dynamic forecasting model. *Applied Energy*. 2021;291:116798. doi:<u>10.1016/j.apenergy.2021.116798</u>

4 Hysolli. Save it in DNA. Wyss Institute. Published August 6, 2019. Accessed March 21, 2023. https://wyss.harvard.edu/news/save-it-in-dna/

5 Banal JL, Shepherd TR, Berleant J, et al. Random access DNA memory using Boolean search in an archival file storage system. *Nat Mater*. 2021;20(9):1272-1280. doi:<u>10.1038/s41563-021-01021-3</u>

6 Chen W, Han M, Zhou J, et al. An artificial chromosome for data storage. *National Science Review*. 2021;8(5):nwab028. doi:<u>10.1093/nsr/nwab028</u>

7 Wilken R, Kennedy J. Everyday data cultures and USB portable flash drives. *International Journal of Cultural Studies*. 2022;25(2):192-209. doi:<u>10.1177/13678779211047917</u>

8 Zheng G, Jing Y. Air Conditioning: Selecting the Optimal Cool Storage System. *Energy & Environment*. 2007;18(2):251-257.

9 Teague R. Ancient DNA and Neanderthals. The Smithsonian Institution’s Human Origins Program. Published September 29, 2022. Accessed March 21, 2023.

10 De Silva PY, Ganegoda GU. New Trends of Digital Data Storage in DNA. *BioMed Research International*. 2016;2016:e8072463. doi:<u>10.1155/2016/8072463</u>

11 Matange K, Tuck JM, Keung AJ. DNA stability: a central design consideration for DNA data storage systems. *Nat Commun*. 2021;12(1):1358. doi:<u>10.1038/s41467-021-21587-5</u>

12 Erlich Y, Zielinski D. DNA Fountain enables a robust and efficient storage architecture. *Science*. 2017;355(6328):950-954. doi:10.1126/science.aaj2038

13 Hughes RA, Ellington AD. Synthetic DNA Synthesis and Assembly: Putting the Synthetic in Synthetic Biology. *Cold Spring Harb Perspect Biol*. 2017;9(1):a023812. doi:<u>10.1101/cshperspect.a023812</u>

14 Matange K, Tuck JM, Keung AJ. DNA stability: a central design consideration for DNA data storage systems. *Nat Commun*. 2021;12(1):1358. doi:<u>10.1038/s41467-021-21587-5</u>

15 Burel A, Carapito C, Lutz JF, Charles L. MS-DECODER: Milliseconds Sequencing of Coded Polymers. *Macromolecules*. 2017;50(20):8290-8296. doi:<u>10.1021/acs.macromol.7b01737</u>

16 Created with BioRender.com.

17 Byron J. CRSS publication: Measuring the Cost of Reliability in Archival Systems. CRSS. Published October 2020. Accessed March 21, 2023. https://www.crss.ucsc.edu/pub/byron-msst20.html

18 Allentoft ME, Collins M, Harker D, et al. The half-life of DNA in bone: measuring decay kinetics in 158 dated fossils. *Proceedings of the Royal Society B: Biological Sciences*. 2012;279(1748):4724-4733. doi:10.1098/rspb.2012.1745

19 Willerslev E, Hansen AJ, Rønn R, et al. Long-term persistence of bacterial DNA. *Current Biology*. 2004;14(1):R9-R10. doi:<u>10.1016/j.cub.2003.12.012</u>

20 Allentoft ME, Collins M, Harker D, et al. The half-life of DNA in bone: measuring decay kinetics in 158 dated fossils. *Proceedings of the Royal Society B: Biological Sciences*. 2012;279(1748):4724-4733. doi:<u>10.1098/rspb.2012.1745</u>

21 Schulz WG, Nieman RA, Skibo EB. Evidence for DNA phosphate backbone alkylation and cleavage by pyrrolo[1,2-a]benzimidazoles: small molecules capable of causing base-pair-specific phosphodiester bond hydrolysis. *Proc Natl Acad Sci U S A*. 1995;92(25):11854-11858.

22 Organick L, Nguyen BH, McAmis R, et al. An Empirical Comparison of Preservation Methods for Synthetic DNA Data Storage. *Small Methods*. 2021;5(5):e2001094. doi:<u>10.1002/smtd.202001094</u>

23 Liu Y, Zheng Z, Gong H, et al. DNA preservation in silk. *Biomater Sci*. 2017;5(7):1279-1292. doi:<u>10.1039/C6BM00741D</u>

24 Kohll AX, Antkowiak PL, Chen WD, et al. Stabilizing synthetic DNA for long-term data storage with earth alkaline salts. *Chem Commun*. 2020;56(25):3613-3616. doi:<u>10.1039/D0CC00222D</u>

25 Chen WD, Kohll AX, Nguyen BH, et al. Combining Data Longevity with High Storage Capacity—Layer-by-Layer DNA Encapsulated in Magnetic Nanoparticles. *Advanced Functional Materials*. 2019;29(28):1901672. doi:<u>10.1002/adfm.201901672</u>

26 Clermont D, Santoni S, Saker S, Gomard M, Gardais E, Bizet C. Assessment of DNA Encapsulation, a New Room-Temperature DNA Storage Method. *Biopreservation and Biobanking*. 2014;12(3):176-183. doi:<u>10.1089/bio.2013.0082</u>

27 Koch J, Gantenbein S, Masania K, Stark WJ, Erlich Y, Grass RN. A DNA-of-things storage architecture to create materials with embedded memory. *Nat Biotechnol*. 2020;38(1):39-43. doi:<u>10.1038/s41587-019-0356-z</u>

28 Grass RN, Heckel R, Puddu M, Paunescu D, Stark WJ. Robust Chemical Preservation of Digital Information on DNA in Silica with Error-Correcting Codes. *Angewandte Chemie International Edition*. 2015;54(8):2552-2555. doi:<u>10.1002/anie.201411378</u>

29 Koch J, Gantenbein S, Masania K, Stark WJ, Erlich Y, Grass RN. A DNA-of-things storage architecture to create materials with embedded memory. *Nat Biotechnol*. 2020;38(1):39-43. doi:<u>10.1038/s41587-019-0356-z</u>

30 Mikutis G, Schmid L, Stark WJ, Grass RN. Length-dependent DNA degradation kinetic model: Decay compensation in DNA tracer concentration measurements. *AIChE Journal*. 2019;65(1):40-48. doi:<u>10.1002/aic.16433</u>

31 Shao W, Khin S, Kopp WC. Characterization of Effect of Repeated Freeze and Thaw Cycles on Stability of Genomic DNA Using Pulsed Field Gel Electrophoresis. *Biopreservation and Biobanking*. 2012;10(1):4-11. doi:<u>10.1089/bio.2011.0016</u>

32 Baoutina A, Bhat S, Partis L, Emslie KR. Storage Stability of Solutions of DNA Standards. *Anal Chem*. 2019;91(19):12268-12274. doi:<u>10.1021/acs.analchem.9b02334</u>

33 Ivanova NV, Kuzmina ML. Protocols for dry DNA storage and shipment at room temperature. *Molecular Ecology Resources*. 2013;13(5):890-898. doi:<u>10.1111/1755-0998.12134</u>

34 Madisen L, Hoar DI, Holroyd CD, Crisp M, Hodes ME, Reynolds JF. The effects of storage of blood and isolated DNA on the integrity of DNA. *American Journal of Medical Genetics*. 1987;27(2):379-390. doi:<u>10.1002/ajmg.1320270216</u>

35 Smith S, Morin PA. Optimal Storage Conditions for Highly Dilute DNA Samples: A Role for Trehalose as a Preserving Agent. *J Forensic Sci*. 2005;50(5):JFS2004411-8. doi:<u>10.1520/JFS2004411</u>

36 Trapmann S, Catalani P, Hoorfar J, Prokisch J, van Iwaarden P, Schimmel H. Development of a novel approach for the production of dried genomic DNA for use as standards for qualitative PCR testing of food-borne pathogens. *Accred Qual Assur*. 2004;9(11):695-699. doi:<u>10.1007/s00769-004-0872-4</u>

37 Dhanasekaran S, Doherty TM, Kenneth J. Comparison of different standards for real-time PCR-based absolute quantification. *Journal of Immunological Methods*. 2010;354(1):34-39. doi:<u>10.1016/j.jim.2010.01.004</u>

38 Newman S, Stephenson AP, Willsey M, et al. High density DNA data storage library via dehydration with digital microfluidic retrieval. *Nat Commun*. 2019;10(1):1706. doi:<u>10.1038/s41467-019-09517-y</u>

39 Bonnet J, Colotte M, Coudy D, et al. Chain and conformation stability of solid-state DNA: implications for room temperature storage. *Nucleic Acids Research*. 2010;38(5):1531-1546. doi:<u>10.1093/nar/gkp1060</u>

40 Johnson CR. Modeling Non-specific Binding in Gel-Based DNA Computers. In: Garzon MH, Yan H, eds. *DNA Computing*. Lecture Notes in Computer Science. Springer; 2008:170-181. doi:<u>10.1007/978-3-540-77962-9_18</u>

41 Bee C, Chen YJ, Ward D, et al. Content-Based Similarity Search in Large-Scale DNA Data Storage Systems. Published online May 27, 2020:2020.05.25.115477. doi:10.1101/2020.05.25.115477

42 Liu K, Song W, Deng Y, et al. Electrooxidation enables highly regioselective dearomative annulation of indole and benzofuran derivatives. *Nature Communications*. 2020;11:3. doi:<u>10.1038/s41467-019-13829-4</u>

43 Lengsfeld CS, Anchordoquy TJ. Shear-Induced Degradation of Plasmid DNA. *JPharmSci*. 2002;91(7):1581-1589. doi:<u>10.1002/jps.10140</u>

44 Lengsfeld CS, Anchordoquy TJ. Shear-Induced Degradation of Plasmid DNA. *JPharmSci*. 2002;91(7):1581-1589. doi:<u>10.1002/jps.10140</u>

45 Wu ML, Freitas SS, Monteiro GA, Prazeres DMF, Santos JAL. Stabilization of naked and condensed plasmid DNA against degradation induced by ultrasounds and high-shear vortices. *Biotechnol Appl Biochem*. 2009;53(Pt 4):237-246. doi:<u>10.1042/BA20080215</u>

46 Levy MS, Collins IJ, Yim SS, et al. Effect of shear on plasmid DNA in solution. *Bioprocess Engineering*. 1999;20(1):7-13. doi:<u>10.1007/s004490050552</u>

47 Organick L, Ang SD, Chen YJ, et al. Random access in large-scale DNA data storage. *Nat Biotechnol*. 2018;36(3):242-248. doi:<u>10.1038/nbt.4079</u>

48 Vinck, A. J. Han. “Coding Concepts and Reed-Solomon Codes.” arXiv, May 2, 2022. https://doi.org/10.48550/arXiv.2205.01044.

49 Press, William H., and John A. Hawkins. “An Indel-Resistant Error-Correcting Code for DNA-Based Information Storage.” arXiv, December 3, 2018. http://arxiv.org/abs/1812.01112.

50 Created with BioRender.com.

51 Ma S, Saaem I, Tian J. Error correction in gene synthesis technology. *Trends in Biotechnology*. 2012;30(3):147-154. doi:<u>10.1016/j.tibtech.2011.10.002</u>

52 Ma S, Saaem I, Tian J. Error correction in gene synthesis technology. *Trends in Biotechnology*. 2012;30(3):147-154. doi:<u>10.1016/j.tibtech.2011.10.002</u>

53 Azevedo RSS, de Sousa JR, Araujo MTF, et al. In situ immune response and mechanisms of cell damage in central nervous system of fatal cases microcephaly by Zika virus. *Sci Rep*. 2018;8(1):1. doi:<u>10.1038/s41598-017-17765-5</u>

54 Gray J, van Ingen C. Empirical Measurements of Disk Failure Rates and Error Rates. Published online January 25, 2007. doi:<u>10.48550/arXiv.cs/0701166</u>

55 Ghemawat S, Gobioff H, Leung ST. The Google File System. In: *Proceedings of the 19th ACM Symposium on Operating Systems Principles*.; 2003:20-43.

56 Patterson DA, Gibson G, Katz RH. A Case for Redundant Arrays of Inexpensive Disks (RAID).

57 Heckel R, Shomorony I, Ramchandran K, Tse DNC. Fundamental limits of DNA storage systems: 2017 IEEE International Symposium on Information Theory, ISIT 2017. *2017 IEEE International Symposium on Information Theory, ISIT 2017*. Published online August 9, 2017:3130-3134. doi:<u>10.1109/ISIT.2017.8007106</u>

58 Shomorony I, Heckel R. Capacity Results for the Noisy Shuffling Channel. Published online February 27, 2019. Accessed March 21, 2023. http://arxiv.org/abs/1902.10832

